# *In vivo* CRISPR screening identifies Fli1 as a transcriptional safeguard that restrains effector CD8 T cell differentiation during infection and cancer

**DOI:** 10.1101/2020.05.20.087379

**Authors:** Zeyu Chen, Eri Arai, Omar Khan, Zhen Zhang, Shin Foong Ngiow, Yuan He, Hua Huang, Sasikanth Manne, Zhendong Cao, Amy E. Baxter, Zhangying Cai, Elizabeth Freilich, Mohammed A. Ali, Josephine R. Giles, Jennifer E. Wu, Allison R. Greenplate, Makoto Kurachi, Kito Nzingha, Viktoriya Ekshyyan, Zhuoyu Wen, Nancy A. Speck, Alexis Battle, Shelley L. Berger, E. John Wherry, Junwei Shi

## Abstract

Improving effector activity of antigen specific T cells is a major goal in cancer immunotherapy. Despite the identification of several effector T cell (T_EFF_)-driving transcription factors (TF), the transcriptional coordination of T_EFF_ biology remains poorly understood. We developed an *in vivo* T cell CRISPR screening platform and identified a novel mechanism restraining T_EFF_ biology through the ETS family TF, Fli1. Genetic deletion of Fli1 enhanced T_EFF_ responses without compromising memory or exhaustion precursors. Fli1 restrained T_EFF_ lineage differentiation by binding to *cis*-regulatory elements of effector-associated genes. Loss of Fli1 increased chromatin accessibility at ETS:RUNX motifs allowing more efficient Runx3-driven T_EFF_ biology. CD8 T cells lacking Fli1 provided substantially better protection against multiple infections and tumors. These data indicate that Fli1 safeguards the developing CD8 T cell transcriptional landscape from excessive ETS:RUNX-driven T_EFF_ cell differentiation. Moreover, genetic deletion of Fli1 improves T_EFF_ differentiation and protective immunity in infections and cancer.

## Introduction

Effective CD8 T cell (T_EFF_) responses are important for control of chronic infections and cancer. Understanding the mechanisms that regulate T_EFF_ cell differentiation is therefore crucial to improving therapeutic approaches for cancer and other diseases. Activation of naïve CD8 T cells (T_N_) during acutely resolved infections or following vaccination results in acquisition of new properties and differentiation into T_EFF_ cells accompanied by dramatic transcriptional and epigenetic remodeling. After antigen clearance, a more terminally differentiated subset of T_EFF_ cells gradually dies off over the ensuing days to weeks, while a small proportion of memory precursors (T_MP_) further differentiates and seeds the long-term memory CD8 T cell (T_MEM_) pool (Kaech and Cui, 2012). During chronic infections and cancer, however, differentiation of responding CD8 T cells is diverted down a path of exhaustion. Under these conditions, T_EFF_ cells become over-stimulated and persist poorly (Angelosanto et al., 2012; Chen et al., 2019b; Khan et al., 2019), whereas a population of activated precursors differentiate into exhausted CD8 T cells (T_EX_)(Chen et al., 2019b; McLane et al., 2019). T_EX_ cells have high expression of multiple inhibitory receptors including PD-1, decreased effector functions compared to T_EFF_ cells, altered homeostatic regulation compared to T_MEM_ cells, and a distinct transcriptional and epigenetic program (Wherry and Kurachi, 2015). Blocking inhibitory receptor pathways like PD-1:PD-L1 can reinvigorate T_EX_ temporarily, restoring proliferative expansion and some effector-like properties (Barber et al., 2006; Huang et al., 2017; Pauken et al., 2016). Such checkpoint blockade has revolutionized cancer treatment, with clinical benefit demonstrated across multiple cancer types (Topalian et al., 2015). Despite of the success of checkpoint blockade, however, most patients do not achieve durable clinical benefit (Sun et al., 2018) and there is a great need to improve the efficacy of these immunotherapies by augmenting T cell differentiation and effector-like activity following checkpoint blockade or during cellular therapies.

There has been considerable interest in defining the populations of T cells responding to checkpoint blockade (Ribas and Wolchok, 2018; Wei et al., 2018) and interrogating the optimal differentiation states for cellular therapies such as CAR T cells (Brown and Mackall, 2019). T_EX_ cells are prominent in human tumors and likely represent a major source of tumor reactive T cells (Duhen et al., 2018; Huang et al., 2017; Simoni et al., 2018). PD-1 pathway blockade mediates clinical benefit, at least in part, due to reinvigoration of T_EX_ cells allowing these cells to re-access parts of the T_EFF_ cell program (Huang et al., 2017; 2019; Pauken et al., 2016). However, limited therapeutic efficacy in many patients is associated with either suboptimal reinvigoration of T_EX_ cells or failure to achieve durable changes in T_EX_ differentiation state (Huang et al., 2017; 2019; Koyama et al., 2016). A key subset of T_EX_ cells required for the therapeutic benefit of PD-1 blockade in mouse models is a progenitor population of T_EX_ cells that, upon PD-1 blockade differentiates into a more numerous terminally exhausted subset (Blackburn et al., 2008). This progenitor subset retains some proliferative capacity, expresses the transcription factor TCF-1 and can be found in both mice (He et al., 2016; Im et al., 2016; Miller et al., 2019; Utzschneider et al., 2016; Wu et al., 2016) and humans (Huang et al., 2019; Sade-Feldman et al., 2018). For cellular therapies, there is evidence that therapeutic failures are associated with the development of exhaustion (Chen et al., 2019a; Fraietta et al., 2018) and approaches that antagonize exhaustion are actively being investigated for CAR T cells (Long et al., 2015; Lynn et al., 2019; Wei et al., 2019). Moreover, cellular therapy products that are more stem T_MEM_-like have improved efficacy (Alizadeh et al., 2019; Sommermeyer et al., 2016). However, a key to both response to checkpoint blockade and cellular therapies to control cancer is the ability to effectively engage a robust effector program, including numerical expansion and elicitation of effector activity. Understanding the underlying molecular mechanisms that control this effector activity is needed to effectively design therapeutic interventions for chronic infections and cancer.

The role of transcription factors (TFs) in regulating gene expression and differentiation of T_EFF_ versus T_MEM_ or T_EX_ has received considerable recent attention (Kaech and Cui, 2012; Wherry and Kurachi, 2015). For example TFs including Batf and Irf4 (Kurachi et al., 2014; Man et al., 2013), T-bet (Intlekofer et al., 2005; Joshi et al., 2007; Kao et al., 2011; Paley et al., 2012; Sullivan et al., 2003), Eomes (Intlekofer et al., 2005; McLane et al., 2013; Paley et al., 2012; Pearce et al., 2003; Zhou et al., 2010), Blimp-1(Kallies et al., 2009; Rutishauser et al., 2009; Shin et al., 2009; Xin et al., 2016), Id-2 (Cannarile et al., 2006; Omilusik et al., 2018; Yang et al., 2011) and others (Kaech and Cui, 2012; McLane et al., 2019) drive T_EFF_ cell differentiation in acute and chronic infections. Batf and Irf4 have an early role in T cell activation and also induce the second wave of transcriptional induction of effector genes (Kurachi et al., 2014; Man et al., 2013). Runx3 induces T_EFF_ gene expression through T-bet and Eomes (Cruz-Guilloty et al., 2009) and also has a role in the early stages of differentiation of tissue resident memory CD8 T cells (T_RM_)(Milner et al., 2017). Most T_EFF_-associated genes and their cognate *cis*-regulatory regions are inaccessible to these TFs in the T_N_ state, therefore, the role of effector-driving TFs must be linked to chromatin accessibility changes that occur during the T_N_ to T_EFF_ transition. Indeed, there is emerging evidence that some of these early operating TFs, such as Batf, may contribute to T_EFF_ gene accessibility through chromatin remodeling (Pham et al., 2019).

In addition to TF that foster T_EFF_ formation, opposing mechanisms must temper complete commitment to effector differentiation. The two alternate cell fates, T_MEM_ and T_EX_, cannot form from fully committed T_EFF_ (Angelosanto et al., 2012; Chen et al., 2019b; Joshi et al., 2007), suggesting that parts of the T_EFF_ program must be antagonized to allow T_MEM_ and T_EX_ to differentiate. The high mobility group (HMG) TF, TCF-1, for example, is essential for development and maintenance of both T_MEM_ and T_EX_ (Chen et al., 2019b; Im et al., 2016; Utzschneider et al., 2016; Wu et al., 2016; Zhou et al., 2010) TCF-1 represses (directly or indirectly) T_EFF_-driving TF such as T-bet and Blimp-1(Tiemessen et al., 2014), and may also have a role in enabling epigenetic changes (Xing et al., 2016). Moreover, a second HMG TF, Tox, is essential for the development of the T_EX_ cell fate by promoting T_EX_ differentiation and repressing of the T_EFF_ lineage differentiation(Alfei et al., 2019; Khan et al., 2019; Scott et al., 2019; Seo et al., 2019; Yao et al., 2019). Tox reprograms the open chromatin landscape, inducing global T_EX_-specific epigenetic changes and repressing chromatin accessibility associated with T_EFF_ differentiation, in part by recruiting epigenetic modifiers including the lysine acetyltransferase Kat7 (Khan et al., 2019). Despite this work, mechanisms that safeguard against commitment to T_EFF_ differentiation remain poorly understood. Such information could be of considerable utility for immunotherapies focused on enhancing anti-tumor or antiviral activity. However, whereas inactivating pathways like TCF-1 or Tox that would de-repress the entire program of T_EFF_ differentiation are of interest, such approaches result in terminal T_EFF_ and may have limited therapeutic benefit because such cells cannot sustain durable responses. Thus, the discovery of mechanisms that selectively de-repress key aspects of T_EFF_ differentiation, particularly those involved in control of numerical expansion and/or protective immunity would be of considerable interest.

Here, we used *in vivo* CRISPR-Cas9 screening in antigen-specific CD8 T cells responding to acute or chronic viral infection to identify key regulators of T_EFF_ and T_EX_. In particular, we were interested in identifying genes that resulted by gain-of-function, improving T_EFF_ differentiation (i.e. an “Up” screen (Kaelin, 2017)). The CRISPR-Cas9 system has been used to interrogate the cancer-immune system through screening in cancer cells (Gerlach et al., 2016; Ishizuka et al., 2019; Manguso et al., 2017; Pan et al., 2018; Wang et al., 2019), *in vitro* in human T cells (Shifrut et al., 2018) or *in vivo* in mouse T cells (Dong et al., 2019; LaFleur et al., 2019b; Shifrut et al., 2018; Wei et al., 2019; Ye et al., 2019). Genome-wide or large pooled screens have identified regulators of T cell responses such as *Dhx37*, a dead-box helicase (Dong et al., 2019) and the phosphatase Ptpn2 (LaFleur et al., 2019b; 2019a). More focused screens for metabolic regulators or membrane proteins have also identified targets such as *Zc3h12a* (enconding REGNASE-1) (Wei et al., 2019) and *Pdia3, Mgat5, Emp1* and *Lag3* (Ye et al., 2019). Many of these targets appear to function by modulating the activity state of the cell through altered signaling or RNA biology. For example, REGNASE-1 is a zinc finger protein that also may regulate mRNA decay (Uehata et al., 2013) and was implicated to function in CD8 T cells through regulation of BATF (Wei et al., 2019). Because Batf has a role in early transcription control of T_EFF_ differentiation, these findings suggest that *in vivo* CRISPR screening could potentially reveal key regulators of differentiation and developmental fate choices. However, the ability to discover fundamental regulators of cellular differentiation state and/or cellular programming via *in vivo* CRISPR/Cas9 screening in CD8 T cells relevant for immunotherapy remains a key goal. Thus, we developed an *in vivo* CRISPR-Cas9 screening platform in primary CD8 T cells focused on central regulators of T cell differentiation and fate decisions. This CD8 T cell CRISPR screening platform used Cas9^+^ antigen specific CD8 T cells combined with an optimized retroviral (RV) based-sgRNA expression strategy (named <u>Op</u>timized <u>T</u> cell In vivo <u>C</u>RISPR <u>S</u>creening system, OpTICS). We focused our screens on TF to identify genes with central regulatory roles in fate decisions and differentiation trajectories in T_EFF_ versus T_EX_ differentiation. Initially, we used the well-established LCMV infection system, in which the biology of T_EFF_ and T_EX_ can be interrogated with high precision, and then extended the findings to other infection and tumor models. This approach identified known key TFs that are essential for T_EFF_ and T_EX_ differentiation including *Batf, Irf4* and *Myc* where loss of function prevented initial T cell activation and differentiation. We identified known TF that repress or restrain T_EFF_ differentiation including *Tcf7, Smad2* and *Tox*. However, this screen also revealed novel central regulators of T_EFF_ differentiation including many that repress optimal differentiation, such as *Irf2, Erg* and *Fli1*, where CRISPR perturbation led to gain-of-function and improved T_EFF_ responses. In particular, we discovered a novel central role for Fli1 in T_EFF_ responses where this TF specifically antagonized the genome-wide function of ETS:RUNX activity and prevented Runx3-driven T_EFF_ biology. Indeed, genetic loss of Fli1 resulted in robustly improved T_EFF_ responses whereas enforced Fli1 expression restrained differentiation. Fli1 prevented chromatin accessibility specifically at ETS:RUNX motifs and loss of Fli1 enabled transcriptional induction of the T_EFF_ program in a Runx3-driven manner. Moreover, loss of Fli1 improved T_EFF_ biology and protective immunity not only during LCMV infection, but also following infection with influenza virus or *Listeria monocytogenes*. Moreover, deletion of Fli1 potently improved anti-tumor immunity. Thus, Fli1 safeguards the developing activated CD8 T cell epigenome from excessive ETS:RUNX-driven T_EFF_ differentiation and disruption of Fli1 activity can be exploited to improve T_EFF_ activity and protective immunity to infections and cancer.

## Results

### Optimized CRISPR-Cas9 for gene editing in murine primary T *in vivo*

Recently, several CRISPR-Cas9-based genetic perturbation approaches have been developed for unbiased genetic screen to reveal novel regulatory pathways in murine primary T cells *in vivo* (Dong et al., 2019; LaFleur et al., 2019b; Wei et al., 2019; Ye et al., 2019). The current methods either adoptive transferring the Cas9-edited bone marrow progenitors into recipient mice which could lead to unexpected developmental effects (LaFleur et al., 2019b) or using genome-wide screening approaches that require large, non-physiological numbers of CD8 T cells *in vivo* (Dong et al., 2019; Ye et al., 2019) which could skew responses to chronic infections and tumors with aberrant effects on disease control and T cell differentiation (Badovinac et al., 2007; 2004; Blackburn et al., 2008; Blattman et al., 2009; Marzo et al., 2005; Wherry et al., 2005). To address these limitations, we set out to develop a system that can 1) perform genetic manipulation in mature CD8 T cells and 2) use a physiological number of cells that mimicking normal T cell development in the setting of viral infections and cancer. To enable gene editing in antigen specific primary CD8 T cells, we crossed LSL-Cas9^+^ mice (Platt et al., 2014) to CD4^CRE+^P14^+^ mice bearing CD8 T cells specific for the LCMV D^b^GP_33-41_ epitope (termed Cas9^+^P14, or C9P14). We expressed the backbone-optimized Cas9 single guide RNA (sgRNA, Grevet et al., 2018) together with a fluorescence-tracking marker in a retroviral (RV) vector (**Figure S1A**). To evaluate gene editing efficiency *in vivo*, we retrovirally transduced C9P14 cells *ex vivo* with either negative control sgRNA (sgCtrl, **Supplementary Table 1**) or sgRNA targeting *Pdcd1* (Encoding PD-1, sgPdcd1, **Supplementary Table 1)** following an optimized murine T cell RV transduction protocol (Kurachi et al., 2017) (**Figure S1B**). The double-positive populations of sgRNA (mCherry^+^) and Cas9 (GFP^+^) CD8 T cells (**Figure S1C**) were adoptively transferred into congenic recipient mice infected with the chronic strain of LCMV (clone13; LCMV-Cl13) (**Figure S1B**). 9 days post infection (p.i.), sgRNA^+^C9P14 cells were isolated for phenotypic and gene editing evaluation. As expected, sgPdcd1 induced antigen specific CD8 T cells expansion 5 fold greater than the control sgRNA (**Figure S1D**), consistent with the genetic knockout of *Pdcd1* (Odorizzi et al., 2015). We found that the sgPdcd1 resulted in both a robust decrease of PD-1 protein level using flow cytometry (**Figure S1E**) and indel mutations in the corresponded genomic locus of *Pdcd1* using sanger-sequencing (**Figure S1F**). Furthermore, we confirmed the high gene editing efficiency of our system by designing sgRNAs targeting *Klrg1* and *Cxcr3* loci (**Figure S1G, Supplementary Table 1**). Collectively, this *in vitro* sgRNA RV transduction in C9P14 followed by *in vivo* adoptive transfer system provides a robust platform to investigate genetic regulatory network of murine CD8 T cells *in vivo*.

### OpTICS enables pooled genetic screening in CD8 T cells *in vivo*

To enable *in vivo* pooled genetic screening in LCMV infection system, we decided to further optimize the parameters of the C9P14 and retroviral sgRNA platform (**Figure 1A**). First, we determined a physiologically relevant number of adoptively transferred CD8 T cells for pooled genetic screening, as the T cell numbers can influence the outcome of infection or tumor models. For example, in the case of LCMV-Cl13 infection model, high numbers of adoptively transferred P14 CD8 T cells can cause lethal immunopathology or convert a chronic infection to an acutely resolved infection (Blattman et al., 2009) preventing the study of T_EX_ cells. We, therefore, limited the maximal number of adoptive transferred CD8 T cells to 1×10^5^ per mouse (approximately 1×10^4^ after take) that was previously shown not to trigger immunopathology or clearance of chronic infection (Chen et al., 2019b; Kao et al., 2011). Next, to evaluate the performance of our system, we performed pooled genetic screening against a focused set of 29 TFs in CD8 T cells *in vivo* using LCMV models. Previously, we found that sgRNA targeting functional important protein domains can substantially improve genetic screening efficiency, as both inframe and frameshift mutations contribute to generating loss-of-function alleles (Shi et al., 2015). We designed and cloned a sgRNA library targeting the DNA-binding domains of 29 TFs and other control genes (e.g. non-selected control sgRNAs and *Pdcd1*) with 4-5 sgRNAs per target. We found that increasing the average input coverage of more than 400 cells per sgRNA after CD8 T cells engraftment could increase the signal to noise ratio, and successfully identify hits, in comparison to 100 cells per sgRNA **(Figure S2A-S2B)**. Third, we assessed the performance of our *in vivo* screen using P14 cells expressing heterogeneous versus homogeneous alleles of the LSL-Cas9 transgene. We observed that Cas9 heterozygous P14 cells outperformed Cas9 homozygous ones in terms of signal-to-noise ratio in different tissues and consistency between independent screens (**Figure S2A-S2C**). We speculated that this is presumably due to reduced off-target DNA damage in the heterozygous setting as we have recently noted for CRE recombinase(Kurachi et al., 2019). From these preliminary optimization screens, we identified *Batf, Irf4*, and *Myc* as essential for early T cell activation because genetic targeting of these genes potently inhibited T cell activation *in vivo* (**Figure S2A-S2B**), consistent with known roles for Batf and Irf4 (Grusdat et al., 2014; Kurachi et al., 2014; Man et al., 2013; 2017; Quigley et al., 2010), and Myc (Lindsten et al., 1988; Wang et al., 2011) in T_EFF_ biology. Of note, this system was highly efficient with up to ∼100-fold enrichment for genes essential for CD8 T cell responses (**Figure S2A-S2B**) and nearly 20-fold enrichment for genes that repress T cell activation and differentiation (**Figure S2D**). Thus, this platform, termed the <u>Op</u>timized <u>T</u> cell In vivo <u>C</u>RISPR <u>S</u>creening system (OpTICS), provides a robust method for focused *in vivo* CRISPR screening in CD8 T cells.

**Figure 1.**
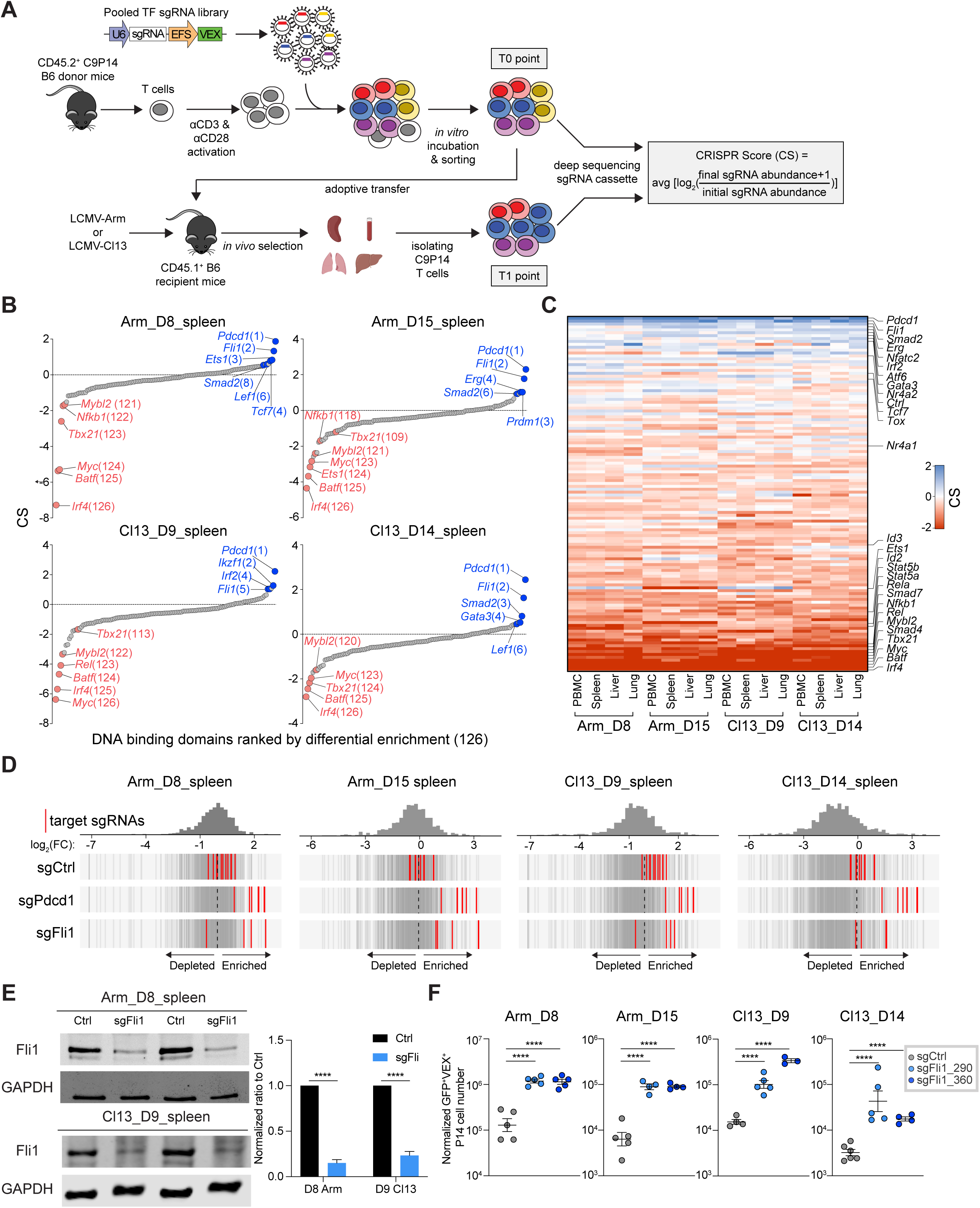
Dissecting transcriptional programs during CD8 T cell response using OpTICS system. A. Experimental design for <u>Op</u>timized <u>T</u> cell In vivo <u>C</u>RISPR <u>S</u>creening (OpTICS). On Day 0(D0), CD8 T cells were isolated from CD45.2^+^ C9P14 donor mice and activated *in vitro*; CD45.1^+^ WT recipient mice were infected with LCMV. On D1 p.i., activated C9P14 cells were transduced with the retroviral (RV) sgRNA library for 6 hours. On D2 p.i., Cas9^+^sgRNA^+^ P14 cells were sort purified; 5-10% of the sorted cells were directly frozen for D2 baseline (T0 time point), and the rest were adoptively transferred into LCMV-infected recipient mice. Cas9^+^sgRNA^+^ P14 cells were then isolated from different organs of recipient mice on the indicated days by MACS and FACS sorting (T1 time point). Genomes were isolated from the Cas9^+^sgRNA^+^ P14 cells isolated at the indicated time points and conditions. Targeted PCR with sequencing adaptors for the sgRNA cassettes was performed and PCR products were sequenced. The CRISPR Score (CS) was calculated according to the formula shown. B. The CS comparing T1 time point to T0 time point (D2 baseline) for different target genes from Cas9^+^sgRNA^+^ cells from spleen on D8 p.i. of Arm, D15 p.i. of Arm, D9 p.i. of Cl13 and D14 p.i. of Cl13. X-axis shows targeted genes; y-axis shows the CS of each targeted gene (calculated using 4-5 sgRNAs). C. Heatmap of CS for targeted genes. Heatmap ranks the geometric means of CS for each targeted gene. Normalization details in Star Methods. D. Distribution of Ctrl, *Pdcd1* and *Fli1* sgRNAs. Axis represents log_2_ fold change (FC). Histogram shows distribution of all sgRNAs. Red bars represent targeted sgRNAs, grey bars represent all other sgRNAs. E. Western blot for Fli1 protein from sorted Cas9^+^sgFli1^+^ P14 cells and paired Cas9^+^sgFli1^-^ P14 cells from spleen. Two *Fli1*-sgRNAs (sgFli1_290 and sgFli1_360) were used. Pooled mice were used in these experiments (3-5 mice for Arm, 10-15 mice for Cl13). Bar graph represents the normalized band intensity of Fli1. The normalized band intensity of Fli1 was first normalized to GAPDH, then a ratio between Cas9^+^sgFli1^+^ versus Cas9^+^sgFli1^-^ was calculated and presented in the bar graph. F. Normalized Cas9^+^sgRNA (VEX)^+^ cell numbers from spleen for Ctrl-sgRNA(sgCtrl) and 2 *Fli1*-sgRNA (sgFli1_290 and sgFli1_360) groups on D8 p.i. of Arm, D15 p.i. of Arm, D9 p.i. of Cl13 and D14 p.i. of Cl13. Cell numbers normalized to the sgCtrl group based on D2 *in vitro* transduction efficiency (see Figure S3B and S3D). \**P*<0.05, \*\**P*<0.01, \*\*\**P*<0.001, \*\*\*\**P*<0.001 versus control (One-Way Anova analysis). Data are representative of 4 independent experiments (mean±s.e.m.) with at least 4 mice/group for F.

### OpTICS identifies novel TF involved in T_EFF_ and T_EX_ cell differentiation

To identify new TFs governing T_EFF_ and T_EX_ cell differentiation, we constructed another domain-focused sgRNA library against 120 TFs (**Supplementary Table 2**, candidate selection in Star Methods). This library has 675 sgRNAs total, including 4-5 sgRNAs per DNA-binding domain, positive selection controls (sgPdcd1), and non-selection controls (e.g. sgAno9, sgRosa26, etc.) (**Figure 1A**). With this 120-TF targeting sgRNA library, we next interrogated *in vivo* selection and sgRNA enrichment at 1 or 2 weeks p.i. with acutely resolving LCMV Arm (LCMV-Arm) or chronic LCMV-Cl13 (**Figure 1A**). We examined C9P14 cells from 4 different anatomical locations (PBMC, spleen, liver and lung) to identify TFs that were broadly important for CD8 T cell responses to infection. In general, groups clustered by time point and infection, rather than anatomical location (**Figure S2E**). At 2 weeks p.i. the data for LCMV-Arm and LCMV-Cl13 diverged (**Figure S2E**), consistent with different trajectories of T cell differentiation during acutely resolved versus chronic infections (Wherry et al., 2007). Focusing on the spleen, *Batf, Irf4* and *Myc* emerged as some of the strongest negative selected hits at both time points in both infections, consistent with the importance of these TFs for initial CD8 T cell activation and T_EFF_ differentiation (**Figure 1B-1C**). We also confirmed several other known effector-driving TFs including *Tbx21* (encoding T-bet), *Id2, Stat5a, Stat5b*, and components of the NF-kB complex (**Figure 1B-1C**). In addition, several TFs with potential novel roles in T cell activation and differentiation were revealed including *Smad4, Smad7* and *Mybl2* (**Figure 1B-1C**).

Many screens are designed to identify loss of function targets such as the TF noted above. In addition to identifying genes necessary for initial CD8 T cell activation and differentiation, we also used the OpTICS system as an “UP” screen (Kaelin, 2017) to identify genes that *repressed* optimal T cell activation and T_EFF_ cell differentiation. Such genes, like *Pdcd1*, represent potential immunotherapy targets for improving efficacy of T cell responses in cancer or infections. PD-1 served as a prototypical positive control where, as expected, *Pdcd1*-sgRNAs were strongly positively selected across infections, time points and in all tissues (**Figure 1B-1D**). This screen also identified TFs that scored as antagonizing robust CD8 T cell responses (**Figure 1B-1C**). Among these, *Tcf7* (encoding TCF-1), *Tox* and *Smad2* have been implicated in limiting robust T_EFF_ cell differentiation, activation and/or expansion. In particular, TCF-1 promotes stem cell-like memory T cells in acutely resolved infections (Zhou et al., 2010) or T_EX_ precursors and progenitors during chronic infections (Chen et al., 2019b; Im et al., 2016; Utzschneider et al., 2016; Wu et al., 2016; Zhou et al., 2010); suppressing TCF-1 has also been shown to augment T_EFF_ cell differentiation (Chen et al., 2019b; Lin et al., 2016). Tox is an essential driver of T_EX_ cell fate and represses terminal T_EFF_ differentiation (Alfei et al., 2019; Khan et al., 2019; Seo et al., 2019; Yao et al., 2019). Smad2 has been shown to limit T_EFF_ cell responses during both acute and chronic infection (Tinoco et al., 2009). We also identified *Nfatc2* and *Nr4a2* (**Figure 1C**), both of which have been implicated in fostering T cell exhaustion and thus limiting T_EFF_ responses (Chen et al., 2019a; Martinez et al., 2015). In addition, *Gata3* identified here has been implicated in driving T cell dysfunction and inhibiting T_EFF_ cell response (Singer et al., 2016). Thus, this screen identified key TFs known to restrain T_EFF_ differentiation and, in some cases, promote exhaustion.

This OpTICS screen also identified novel TFs that restrained optimal T_EFF_ differentiation. This set of genes included *Atf6, Irf2, Erg* and *Fli1*, with *Fli1* among the strongest hits in repressing T_EFF_ cell differentiation. The identification of Fli1 as a repressor of T_EFF_ differentiation occurred similarly in LCMV-Arm and LCMV-Cl13 infections indicating a common role for this TF in restraining T_EFF_ biology in settings of acutely resolving or chronic infection. In both infections, *Fli1*-sgRNAs were the top ranking positively selected sgRNAs across different conditions (**Figure 1B-1D**). We therefore focused additional efforts on interrogating the biology of Fli1 in CD8 T cell differentiation.

Fli1 is an ETS family TF with roles in hematopoiesis and other developmental pathways in non-immune cell types (Kruse et al., 2009; Pimanda et al., 2007; Tijssen et al., 2011). In hematopoiesis, Fli1 has been implicated in regulating stem cell maintenance and differentiation (Cai et al., 2015). The role of Fli1 in T_EFF_, T_MEM_ or T_EX_ cells, however, is unknown. To interrogate this question, we selected 2 *Fli1*-sgRNAs (sgFli1_290 and sgFli1_360) and confirmed that these sgRNAs effectively edited the *Fli1* gene (70%-80% editing; **Figure S3A**) leading to reduced protein expression (**Figure 1E**). Targeting Fli1 using these individual *Fli1-*sgRNAs in C9P14 cells *in vivo* resulted in 5-20 fold greater expansion at 1 and 2 weeks p.i. with either LCMV-Arm or LCMV-Cl13 (**Figure 1F** and **S3B-S3E**). These data indicate repression of robust CD8 T cell expansion by Fli1 in acutely resolving or developing chronic infection.

### Genetic deletion of Fli1 promotes robust T_EFF_-differentiation during acutely resolved infection

We next interrogated the differentiation state of *Fli1*-sgRNA (sgFli1) or Ctrl-sgRNA (sgCtrl) transduced C9P14 cells during acutely resolved infection. On Day 8 p.i., Fli1 deletion reduced the proportion of the CD127^Hi^ memory precursors (T_MP_), whereas the frequency KLRG1^Hi^ terminal effector (T_EFF_) population remained unchanged at this time point consistent with a slight increase in the CD127^Lo^KLRG1^Lo^ population (**Figure 2A**). These effects were more dramatic at Day 15 p.i. resulting in a reduction in the CD127^Hi^ T_MP_ population and, at this time point, an increase in the proportion of KLRG1^Hi^ T_EFF_ cells (**Figure 2A**). However, at both time points the absolute number of both T_MP_ and T_EFF_ was increased due to the proliferative expansion of *Fli1*-deficient CD8 T cells (**Figure 2A**). This skewing of T cell differentiation towards T_EFF_-like populations was also observed when CX3CR1 and CXCR3 were used to identify T_MEM_- and T_EFF_-like cells (Gerlach et al., 2016), with sgFli1 significantly enriching for the CX3CR1^+^CXCR3^-^ T_EFF_ population compared to the CX3CR1^-^CXCR3^+^ early T_MEM_ subset (**Figure 2B**). To further interrogate the causality of Fli1 in these effects on CD8 T cell differentiation, we next enforced Fli1 expression in WT LCMV-specific P14 cells using a retroviral (RV) based overexpression (OE) system (Kurachi et al., 2017). A ∼5-fold reduction in responding Fli1-OE-RV transduced P14 cells was observed Day 8 and Day 16 p.i. compared to the empty vector control (**Figure 2C**). Furthermore, enforced Fli1 expression also skewed responding P14 cells towards T_MP_ differentiation, with an increase in CD127^Hi^KLRG1^Lo^ and CX3CR1^-^CXCR3^+^ populations at the expense of KLRG1^Hi^CD127^Lo^ or CX3CR1^+^CXCR3^-^ T_EFF_ cells (**Figure 2D-2E**). Together, these data reveal a role for Fli1 in restraining T_EFF_ differentiation and promoting T_MP_ development during acute infection.

**Figure 2.**
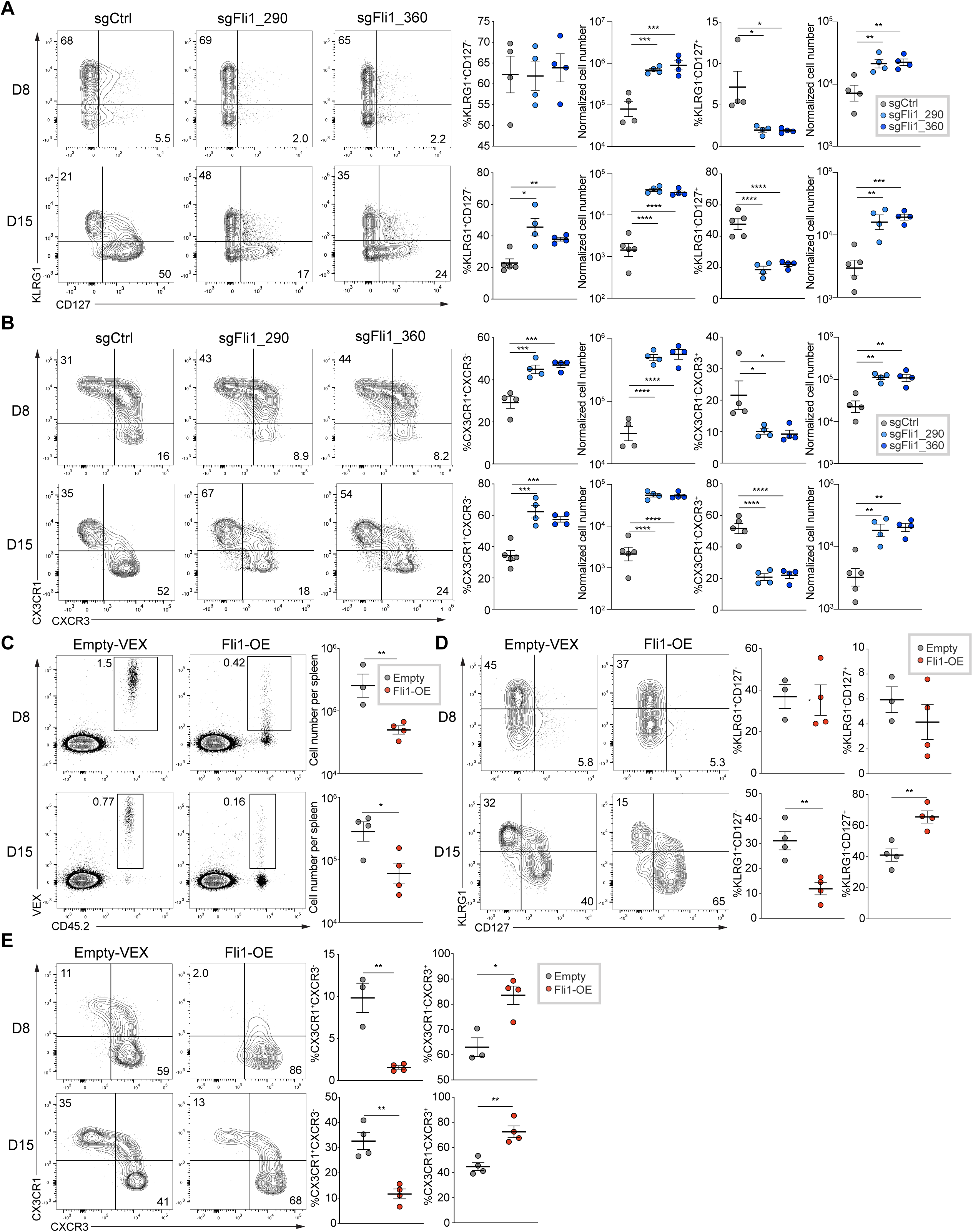
Fli1 inhibits T_EFF_ cell proliferation and differentiation during acute infection. A. Flow cytometry plots and statistical analysis of KLRG1^Hi^CD127^Lo^ Terminal Effectors (TEs) and KLRG1^Lo^CD127^Hi^ Memory Precursors (MPs). Frequencies (left) and numbers (right) from spleen for sgCtrl and 2 sgFli1 groups on D8 and D15 p.i. of Arm. Gated on Cas9(GFP)^+^ sgRNA(VEX)^+^ P14 cells. B. Flow cytometry plots and statistical analysis of CX3CR1^+^CXCR3^-^ T_EFF_ cells and CX3CR1^-^CXCR3^+^ early T_MEM_ cells frequencies (left) and numbers (right) from spleen for sgCtrl and 2 sgFli1 groups on D8 and D15 p.i. of Arm. Gated on Cas9^+^sgRNA^+^ P14 cells. (C-E) Experimental design. On D0, CD45.1^+^ P14 cells were activated and recipient mice were infected with Arm; On D1 p.i., activated P14 cells were transduced with Empty-RV or Fli1-over expression(OE)-RV for 6 hours. On D2 p.i., VEX^+^ P14 cells were purified by sorting from Empty-RV or Fli1-OE-RV groups, and 5 x 10^4^ cells were adoptively transferred into the infected recipient mice. C. Flow cytometry plots of CD45.2^+^VEX^+^ cell frequency and statistical analysis of CD45.2^+^VEX^+^ cell number for Empty-RV and Fli1-OE-RV conditions. D. Flow cytometry plots and statistical analysis of KLRG1^Hi^CD127^Lo^ TE and KLRG1^Lo^CD127^Hi^ MP frequencies from spleen for Empty-RV and Fli1-OE-RV groups on D8 and D15 p.i. of Arm. Gated on VEX^+^ P14 cells. E. Flow cytometry plots and statistical analysis of CX3CR1^+^CXCR3^-^ T_EFF_ cell and CX3CR1^-^CXCR3^+^ early T_MEM_ cell frequencies from spleen for Empty-RV and Fli1-OE-RV groups on D8 and D15 p.i. of Arm. Gated on VEX^+^ P14 cells. \**P*<0.05, \*\**P*<0.01, \*\*\**P*<0.001, \*\*\*\**P*<0.001 versus control (two-tailed Student’s *t*-test and One-Way Anova analysis). Data are representative of 2-4 independent experiments (mean±s.e.m.) with at least 3 mice/group.

### Fli1 antagonizes T_EFF_-like differentiation during chronic infection

During chronic viral infection, there is an early fate bifurcation for CD8 T cell responses where antiviral CD8 T cells develop into either T_EFF_-like cells or form T_EX_ precursors that ultimately seed the mature T_EX_ population (Chen et al., 2019b; Khan et al., 2019). We therefore investigated the role of Fli1 in this cell fate decision during the first 1-2 weeks of chronic infection. As in acutely resolving infection, genetic perturbation of Fli1 resulted in a skewing of the virus-specific CD8 T cell response towards the T_EFF_ pathway, as defined by TCF-1^-^GrzmB^+^ or Ly108^-^CD39^+^ (**Figure 3A** and (Chen et al., 2019b)). Due to the 5-10-fold increase in total *Fli1*-deficient cells, the cell numbers of both the T_EFF_-like and T_EX_ precursor populations were increased (**Figure S4A-S4B**). In contrast, enforced Fli1 expression not only resulted in lower cell numbers (**Figure S4C**), but also fostered formation of Ly108^+^CD39^-^ or TCF-1^+^GrzmB^-^ T_EX_ precursors (**Figure S4D-S4E**). Moreover, genetic perturbation of Fli1 resulted in an increased proportion of T_EFF_-like cells marked by high CX3CR1 or Tim3 expression (**Figure 3B** and (Chen et al., 2019b; Zander et al., 2019)). These data were confirmed by enforced Fli1 expression during LCMV-Cl13 infection that again provoked the opposite effect (**Figure S4F**). Notably, PD-1 expression was unchanged and, unlike acutely resolved infection, KLRG1 expression was unaffected (**Figure 3B**).

**Figure 3.**
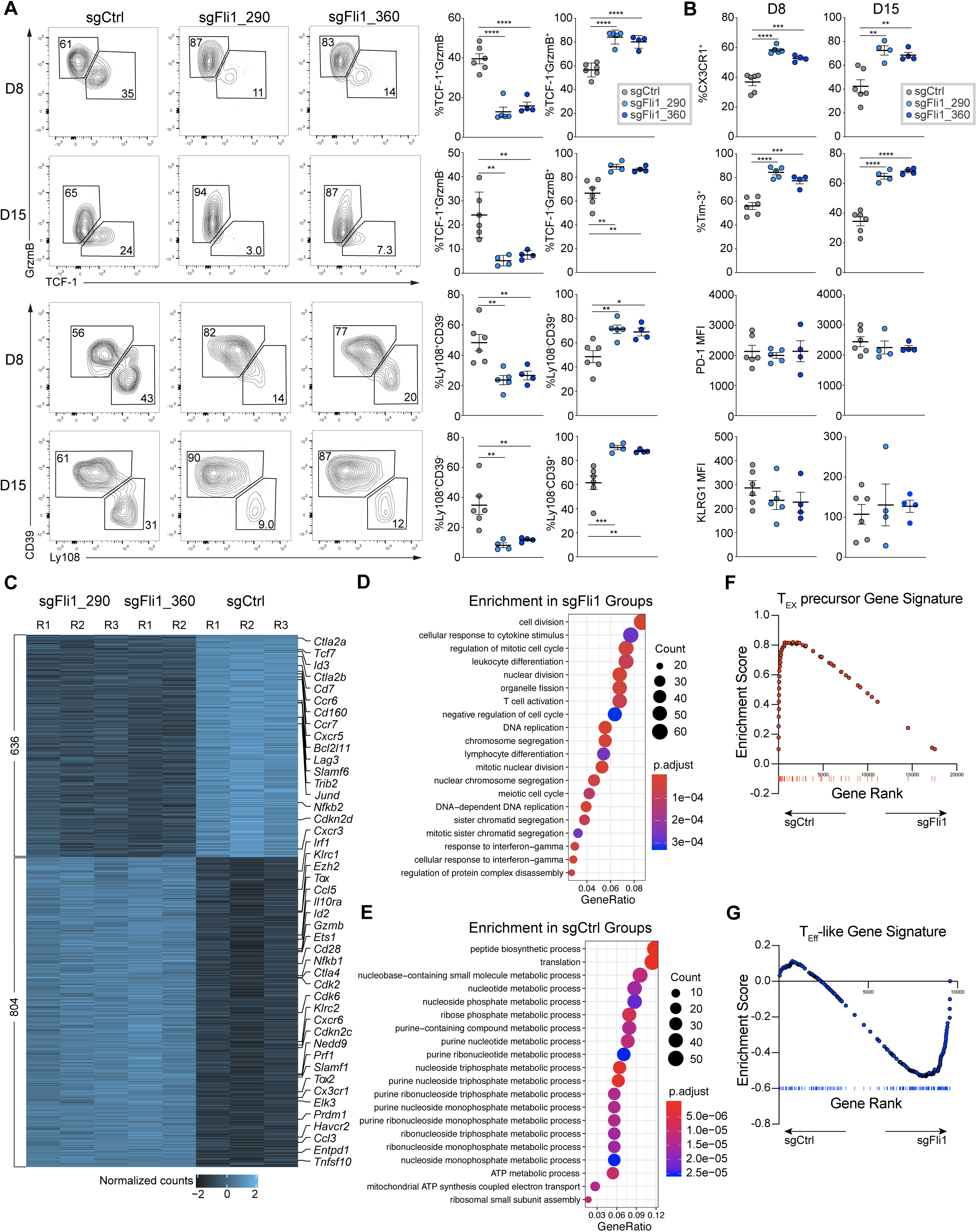
Fli1 antagonizes T_EFF_-like cell differentiation during chronic infection. A. Flow cytometry plots and statistical analysis of Ly108^-^CD39^+^ or TCF-1^-^Gzmb^+^ T_EFF_-like cells and Ly108^+^CD39^-^ or TCF-1^+^Gzmb^-^ T_EX_ precursor frequencies from spleen for sgCtrl and 2 sgFli1 groups on D8 and D15 p.i. of Cl13. Gated on the Cas9(GFP)^+^sgRNA(VEX)^+^ P14 cells. B. Statistical analysis of CX3CR1^+^ and Tim-3^+^ frequencies, and KLRG1 and PD-1 MFIs from spleen for sgCtrl and 2 sgFli1 groups on D8 and D15 p.i. of Cl13. Gated on the Cas9^+^sgRNA^+^ P14 cells. C. Heatmap showing differentially expressed genes between sgCtrl and 2 sgFli1 groups. D. Gene Ontology (GO) enrichment analysis for the sgFli1 groups. E. GO enrichment analysis for the sgCtrl groups. F. Gene Set Enrichment Analysis (GSEA) of T_EX_ precursor signature (adapted from Chen et al. 2019) between sgCtrl and sgFli1 groups. G. GSEA of T_EFF_-like signature (adapted from Bengsch et al. 2018) between sgCtrl and sgFli1 groups. \**P*<0.05, \*\**P*<0.01, \*\*\**P*<0.001, \*\*\*\**P*<0.001 versus control (two-tailed Student’s *t*-test and One-Way Anova analysis). Data are representative of 4 independent experiments (mean±s.e.m.) with at least 4 mice/group for A and B.

To dissect the underlying mechanism by which Fli1 regulates CD8 T cell differentiation, we performed RNA-seq on sorted sgCtrl^+^ or sgFli1^+^ C9P14 cells on Day 9 of LCMV-Cl13 infection. Both sgRNAs targeting *Fli1* resulted in a similar transcription effect (**Figure 3C, S4G**). In contrast, the sgFli1^+^ C9P14 cells were transcriptionally distinct from the sgCtrl^+^ C9P14 cells with over 1400 genes differentially expressed between the two conditions (**Figure 3C, S4G**). Among the major changes were a robust increase in effector-associated genes such as *Prf1, Gzmb, Cd28, Ccl3* and *Prdm1* in the sgFli1^+^ C9P14 cells, whereas the sgCtrl^+^ C9P14s were enriched in T_EX_ precursor genes such as *Tcf7, Cxcr5, Slamf6* and *Id3* (**Figure 3C**). Gene Ontology enrichment analysis also identified cell division-associated and T cell activation-associated pathways in sgFli1^+^ C9P14 cells (**Figure 3D**), whereas sgCtrl^+^ C9P14s enriched in multiple metabolic pathways, in particular nucleotide, nucleoside and purine biosynthesis (**Figure 3E**). We next used gene set enrichment analysis (GSEA) to examine skewing between signatures of subpopulations during the early phase of chronic infection, when a divergence of differentiation into either T_EFF_-like or T_EX_ precursor cell fates occurs. Indeed, the T_EX_ precursor signature (Chen et al., 2019b) was strongly enriched in sgCtrl^+^ C9P14 cells compared to the sgFli1^+^ C9P14 population whereas a T_EFF_ gene signature (Bengsch et al., 2018) was strongly enriched in the sgFli1^+^ C9P14 population (**Figure 3F-3G**). Thus, Fli1 repressed optimal T_EFF_ differentiation in both acutely resolving and chronic infection and loss of Fli1 antagonized the development of T_EX_ cells. However, although genetic perturbation of *Fli1* drove an increase in expression of effector-associated genes at Day 9 of chronic infection, *Tox, Tox2* and *Cd28* were also increased. This effect may suggest that although loss of Fli1 might enhance T_EFF_-like biology and might not be at the expense of genes necessary to sustain responses in chronic infection of cancer.

### Fli1 remodels the epigenetic landscape of CD8 T cells and antagonizes T_EFF_-associated gene expression

In acute myeloid leukemia, FLI1 co-localizes with the chromatin remodeler BRD4 (Roe et al., 2015). Moreover, the oncogenic EWS-FLI1 fusion protein can bind genomic microsatellites (Gangwal et al., 2008), trigger *de novo* enhancer formation via chromatin remodeling and inactivate existing enhancers by displacing ETS family members (Riggi et al., 2014). Though EWS-FLI1 and FLI1 have similar GGAA-binding motifs (Riggi et al., 2014; Roe et al., 2015), the prion-like domain from EWS changes the overall function of the EWS-FLI1 fusion likely leading to distinct mechanisms of action (Boulay et al., 2017). Thus, it is unclear how Fli1 affects epigenetic landscape changes in general, especially in developing T_EFF_, T_MEM_ or T_EX_ cells.

To examine the role of Fli1 in supporting the epigenetic landscape of CD8 T cells, we performed ATAC-seq on sgFli1^+^ and sgCtrl^+^ C9P14 cells on Day 9 p.i. with LCMV-Cl13. Compared to sgCtrl^+^ C9P14 cells, the sgFli1^+^ group had considerable changes in chromatin accessibility (**Figure 4A**). Over 5000 open chromatin regions (OCRs) differed between the control and *Fli1-*perturbed groups with approximately equal numbers of peaks gained or lost (**Figure 4B-4D**). Most of these changes were located in intronic or intergenic regions consistent with *cis*-regulatory or enhancer elements (**Figure 4C**).

**Figure 4.**
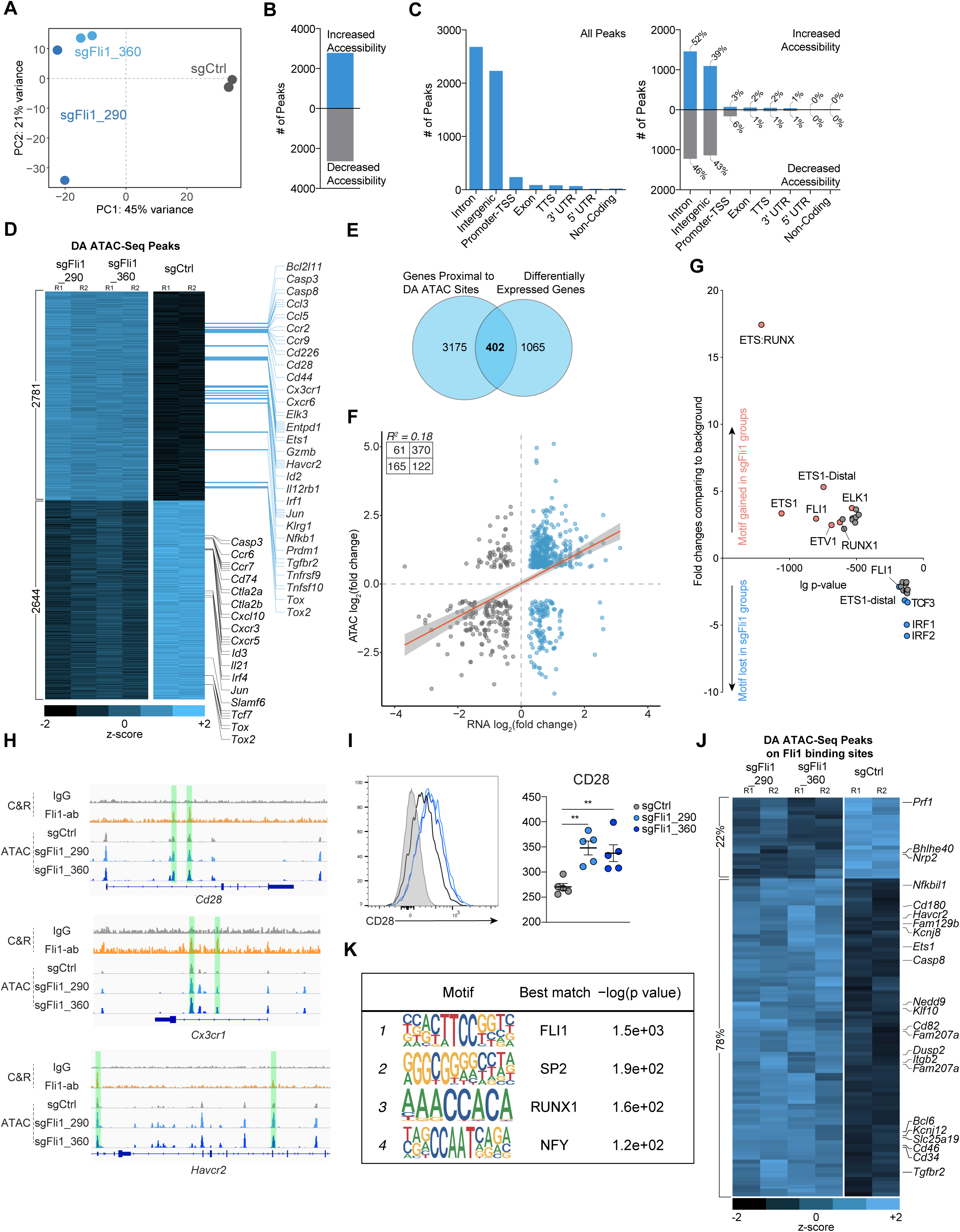
Fli1 reshapes the epigenetic profile of CD8 T cells and inhibits T_EFF_-associated gene expression. A. PCA plot of ATAC-seq data for sgCtrl, sgFli1_290 and sgFli1_360 groups on D9 p.i. of Cl13. B. Overall open chromatin region (OCR) peak changes for sgFli1 groups compared to sgCtrl group. C. Categories of *Cis*-element OCR peaks that changed between sgCtrl and sgFli1 groups. Left plot represents all changes; right plot represents changes for increased or decreased accessibility. D. Heatmap shows differentially accessible peaks between sgCtrl group and 2 sgFli1 groups (adjusted p-value<0.05, Log_10_ Fold Change>0.6). Some of the nearest genes assigned to the peaks are listed. E. Overlapping Venn plot of the genes with differentially accessible (DA) peaks and differentially expressed genes from Figure 3C. F. Pearson correlation of the peak accessibility of the nearest genes versus the differential expression of the genes. G. Transcription factor (TF) motif gain or loss associated with loss of Fli1. X-axis represents the logP-value of the motif enrichment. Y-axis represents the fold change of the motif enrichment. Targeted motifs in the changed OCR between the sgCtrl and sgFli1 groups were compared to the whole genome background to calculate p-value and fold change. H. IgG or Fli1 binding signals from CUT&RUN on P14 cells on D8 p.i. of Cl13 and OCR signals detected by ATAC-seq for sgCtrl-sgRNA, sgFli1_290 and sgFli1_360 groups at the *Cd28, Cx3cr1*, and *Havcr2* loci. I. Histogram of CD28 staining and statistical analysis for sgCtrl, sgFli1_290 and sgFli1_360 groups on D8 Cl13 p.i. Grey shows CD28 staining of CD44^-^ naïve T cell. J. Heatmap shown differentially accessible (DA) peaks overlapped with the Fli1 CUT&RUN binding peaks between sgCtrl group and 2 sgFli1 groups. Some of the nearest genes assigned to the peaks are listed. K. Top 4 enriched TF motifs in the Fli1 CUT&RUN peaks are listed. \**P*<0.05, \*\**P*<0.01 versus control (One-Way Anova analysis). Data are representative of 2 independent experiments (mean±s.e.m.) with at least 5 mice/group for I.

To define how these changes in OCRs could impact the differentiation of T_EFF_ or T_EX_ cells, we next assigned each OCR to the nearest gene to estimate genes that could be regulated by these *cis*-regulatory elements. T_EFF_-associated genes, such as *Ccl3, Ccl5, Cd28, Cx3cr1* and *Prdm1*, gained chromatin accessibility in the sgFli1^+^ group (**Figure 4D**), consistent with the RNA-seq data. In contrast, there was decreased chromatin accessibility near genes involved in T cell progenitor biology such as *Tcf7, Slamf6, Id3* and *Cxcr5* (**Figure 4D**) in the sgFli1^+^ group. Moreover, the sgFli1^+^C9P14 cells had altered accessibility in the *Tox* (and *Tox2)* locus, but these changes included both increased and decreased accessibility across different peaks (**Figure 4D**), consistent with the complex transcriptional and epigenetic control of *Tox* expression (Khan et al., 2019). These changes in chromatin accessibility corresponded to changes in gene expression with ∼1/3 of the genes that changed transcriptionally associated with a differentially accessible chromatin region (402 out of 1467) (**Figure 4E**). In general, increased accessibility correlated with increased transcription though there was a clear subset of regions where decreased accessibility corresponded to increased transcription suggesting potential negative regulatory elements controlled by Fli1 (**Figure 4F**).

To define the mechanisms by which these Fli1-mediated chromatin accessibility changes regulated T cell differentiation, we next defined the TF motifs present in the OCRs that were dependent on Fli1 for altered accessibility. Among the OCRs that decreased in accessibility in the absence of Fli1, the most enriched TF motifs were for IRF1 and IRF2 (**Figure 4G**), potentially linking Fli1 activity to the role of IRF1 and IRF2 downstream of IFN signaling or to the regulation of cell cycle by these TFs (Choo et al., 2006). In the group of OCRs that increased in accessibility in the absence of Fli1, ETS and RUNX motifs were highly enriched (**Figure 4G**). In particular, the composite ETS:RUNX motif was by far the most changed with 18-fold enrichment (**Figure 4G**). These observations suggested that Fli1 may limit the activity of other ETS family members (e.g. ETS1, ETV1 or ELK1) or alter accessibility at ETS:RUNX binding sites (**Figure 4G**). Runx3 is a central driver of T_EFF_ differentiation and functions by directly regulating effector gene expression, coordinating and enabling the effector gene regulation via T-bet and Eomes and antagonizing TCF-1 expression (Cruz-Guilloty et al., 2009; Shan et al., 2017; Wang et al., 2018). Thus, a potential role for Fli1 in Runx3 biology would provide a mechanistic link between loss of Fli1 and improved T_EFF_ differentiation.

We next tested how Fli1 genomic binding was related to changes in chromatin accessibility and T_EFF_ biology, using Fli1 CUT&RUN (Skene et al. 2017). At Day 9 p.i. with LCMV-Cl13 >90% of the identified Fli1 binding sites were contained in OCRs detected by ATAC-Seq (**Figure S4H-S4I**). Specifically, Fli1 bound to OCRs of T_EFF_-like genes such as *Cx3cr1, Cd28* and *Havcr2.* Chromatin accessibility increased at these locations upon Fli1 deletion (**Figure 4H**), resulting in increased transcription (**Figure 3C**) and protein expression (**Figure 3B** and **4I**). In contrast, for genes involved in progenitor biology that were decreased in expression in the absence of Fli1 such as *Tcf7* and *Id3*, direct binding of Fli1 was not observed (**Figure S4H**), likely indicating that the major role of Fli1 is to safeguard against an overly robust T_EFF_ program rather directly enabling memory/progenitor biology. Furthermore, 78% of the sites where Fli1 was defined to bind by CUT&RUN increased in chromatin accessibility in the absence of Fli1; in contrast, 22% decreased in accessibility (**Figure 4J**), suggesting that Fli1 mainly functions to repress chromatin accessibility. Analyzing the DNA binding motifs in the Fli1 CUT&RUN data revealed the expected FLI1 motif. However, SP2, NFY1 and RUNX1 motifs were also significantly enriched where Fli1 bound (**Figure 4K**). Together with the increase in ETS:RUNX motifs in the *Fli1*-deficient ATAC-seq data above, these data support a model where Fli1 coordinates with RUNX family members to control T_EFF_ differentiation.

### Enforced Runx3 expression synergizes with *Fli1*-deletion to enhance T_EFF_ responses

RUNX family TFs are key regulators of CD8 T cell biology. In particularly, Runx3 is central to the transcriptional cascade that coordinates T_EFF_ differentiation (Cruz-Guilloty et al., 2009; Wang et al., 2018). Runx3 also antagonizes a follicular-like CD8 T cell fate by inhibiting TCF-1 expression (Shan et al., 2017) and fosters the development of tissue and tumor resident CD8 T cells (Milner et al., 2017). Runx1, in contrast, is antagonized by Runx3 during T_EFF_ differentiation (Cruz-Guilloty et al., 2009). The roles of Runx1 and Runx3 in T_EX_ development in the early phase of chronic infection are less clear. Because T_EFF_ and T_EX_ are opposing fates in chronic infection (Chen et al., 2019b; Khan et al., 2019; Yao et al., 2019; Zander et al., 2019) and ETS:RUNX motifs become more accessible in the absence of Fli1, we hypothesized that a RUNX-Fli1 axis might influence T_EFF_ versus T_EX_ differentiation. We tested whether Runx1 or Runx3 expression in *Fli1*-deficient CD8 T cells would impact T_EFF_ differentiation in early chronic infection.

Enforced expression of Runx1 in WT P14 cells reduced cell numbers at Day 7 p.i. with LCMV-Cl13 infection (**Figure S5A**) suggesting that Runx1 antagonized CD8 T cell activation and differentiation. Moreover, Runx1-OE fostered formation of Ly108^+^CD39^-^ T_EX_ precursors at the expense of the more T_EFF_-like Ly108^-^CD39^+^ population at this early time point (**Figure S5A**). Next, to interrogate the impact of enforced Runx1 expression in the absence of Fli1, we used a dual RV transduction approach and combined control or Fli1 sgRNA RV transduction with either empty or Runx1 expressing RVs. Singly versus dually transduced cells were distinguished using VEX (for sgRNA) and mCherry. C9P14 cells were transduced and adoptively transferred into LCMV-Cl13-infected mice and the double transduced (i.e. GFP^+^VEX^+^mCherry^+^) C9P14 cells were analyzed at Day 8 p.i. (**Figure 5A**). Dual-transduction was efficient and allowed robust detection of GFP^+^VEX^+^mCherry^+^ C9P14 cells (**Figure S5B**). In the sgFli1+Runx1-OE group the number of GFP^+^VEX^+^mCherry^+^ C9P14 cells was reduced and there were fewer Ly108^-^CD39^+^ cells. In contrast the sgFli1+Empty-RV group had an increase in the GFP^+^VEX^+^mCherry^+^ C9P14 cell population and these cells were skewed towards the Ly108^-^CD39^+^ T_EFF_-like fate (**Figure 5B-5D, S5C**) as above. However, in the *Fli1*-deficient setting where there is enhanced CD8 T cell expansion, Runx1 overexpression reduced the magnitude of the response and partially reversed the skewing towards the Ly108^-^CD39^+^ T_EFF_-like fate caused by loss of Fli1 (**Figure 5B-5D**).

**Figure 5:**
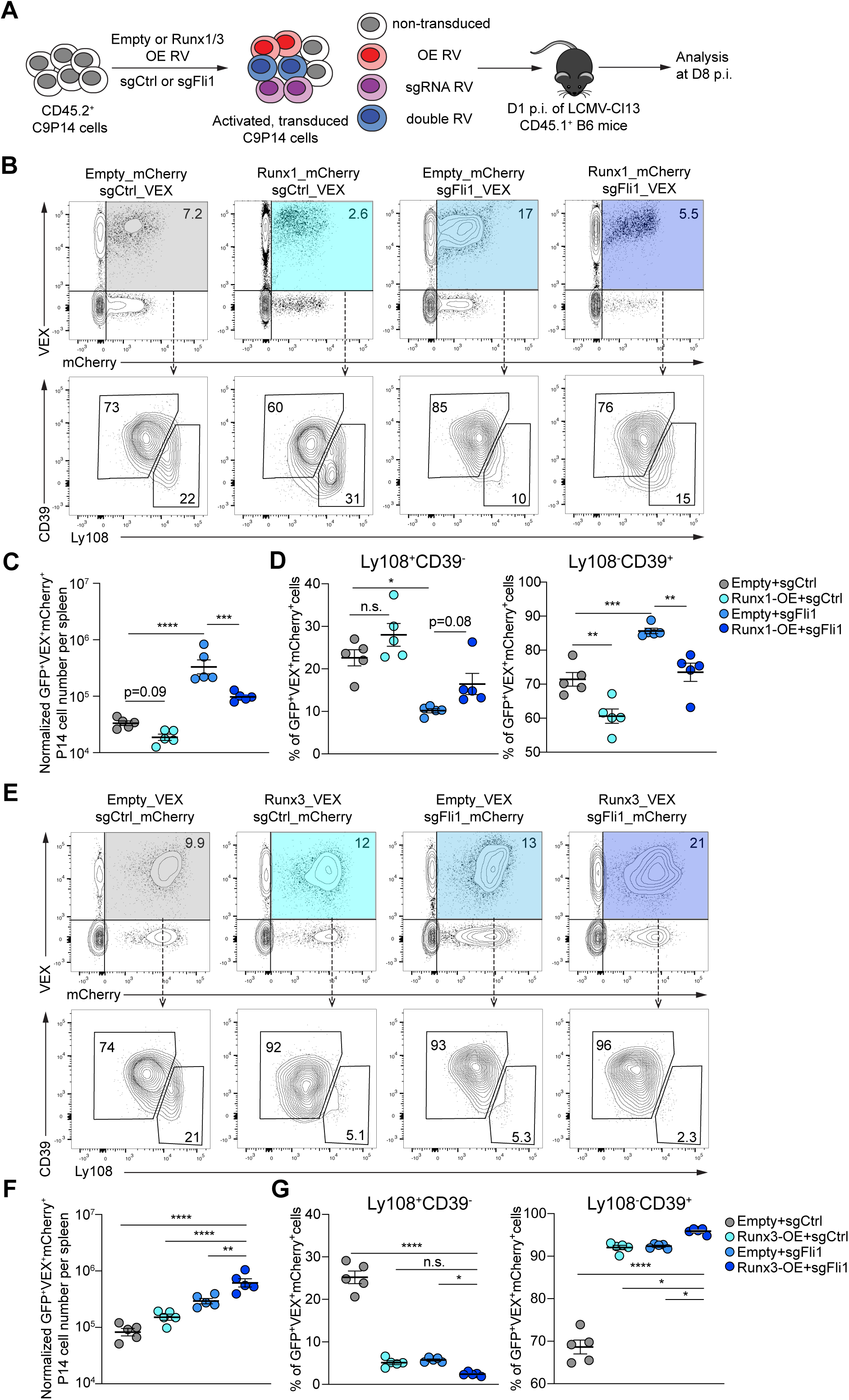
Overexpression of Runx1 inhibits the phenotypes of *Fli1*-deficiency while overexpression of Runx3 enhances *Fli1*-deficient effects. A. Experimental design. On D0, CD8 T cells were isolated from CD45.2^+^ C9P14 donor mice and activated; CD45.1^+^ WT recipient mice were infected with LCMV Cl13. On D1 p.i., activated C9P14 cells were transduced with sgRNA-RV or OE-RV for 12 hours, and 1 x 10^5^ transduced cells were adoptively transferred into infected recipient mice. B-D. Flow cytometry plots (B) and statistical analysis (C-D) of VEX^+^mCherry^+^ C9P14 cells and Ly108^-^CD39^+^/Ly108^+^CD39^-^ C9P14 cells from spleen for sgCtrl-VEX + Empty-mCherry, sgCtrl-VEX + Runx1-mCherry, sgFli1_290-VEX + Empty-mCherry and sgFli1_290-VEX + Runx1-mCherry on D8 p.i. of Cl13. Gated on Cas9(GFP)^+^CD45.2^+^ P14 cells. E-G. Flow cytometry plots (E) and statistical analysis (F-G) of VEX^+^mCherry^+^ C9P14 cells and Ly108^-^CD39^+^/Ly108^+^CD39^-^ C9P14 cells from spleen for sgCtrl-mCherry + Empty-VEX, sgCtrl-mCherry + Runx3-VEX, sgFli1_290-mCherry + Empty-VEX and sgFli1_290-mCherry + Runx3-VEX at D8 p.i. of Cl13. Gated on Cas9^+^CD45.2^+^ P14 cells. \**P*<0.05, \*\**P*<0.01, \*\*\**P*<0.001, \*\*\*\**P*<0.001 versus control (One-Way Anova analysis). Data are representative of 2 independent experiments (mean±s.e.m.) with at least 5 mice/group.

We next investigated the impact of enforced expression of Runx3 in the absence of Fli1 using the same dual transduction approach (**Figure 5A, 5E-5G, S5D-S5E**). Enforced expression of Runx3 alone (in the sgCtrl^+^ group), modestly increased the magnitude of the CD8 T cell response but robustly skewed the GFP^+^VEX^+^mCherry^+^ C9P14 population towards a CD39^+^Ly108^-^ T_EFF_-like population (**Figure 5E-5G**). These effects were more dramatic in the absence of Fli1 with greater numerical amplification and even further skewing towards CD39^+^Ly108^-^ T_EFF_-like cells in the sgFli1+Runx3-OE enforced expression group in both proportion and cell number (**Figure 5E-5G**). While the sgFli1+Runx3-OE group shows reducing the frequency of the T_EX_ precursor population, this group did not show a cell number reduction of this population (**Figure 5E** and **5G**; **S5E**). Taken together, these data support a model where loss of Fli1 reveals ETS:RUNX motifs that can be used by Runx1 and/or Runx3. However, whereas Runx3 drives a more T_EFF_-like population, an effect amplified in the absence of *Fli1*, Runx1 appears to antagonize T_EFF_ generation, in agreement with the opposing functions of Runx1 and Runx3 (Cruz-Guilloty et al., 2009). Thus, Fli1 restrains the T_EFF_ promoting activity of Runx3 function by restricting genome access and protecting ETS:RUNX binding sites. These data reveal Fli1, Runx3 and perhaps Runx1 (as a modest Runx3 antagonist) as key regulators of the fate choice between T_EFF_ and T_EX_ early after initial activation.

### Augmented T_EFF_ cell responses in the absence of Fli1 improve protective immunity against pathogens

The data above provoke the question of whether loss of Fli1 would improve control of infections due to the augmented T_EFF_ differentiation. To test this idea, we used LCMV-Cl13 to investigate protective immunity during chronic infection and two models of acute infection with influenza virus (PR8) or *Listeria monocytogenes* (LM) each expressing the LCMV GP_33-41_ epitope (PR8_GP33_ and LM_GP33_) recognized by P14 cells (**Figure 6A**).

**Figure 6.**
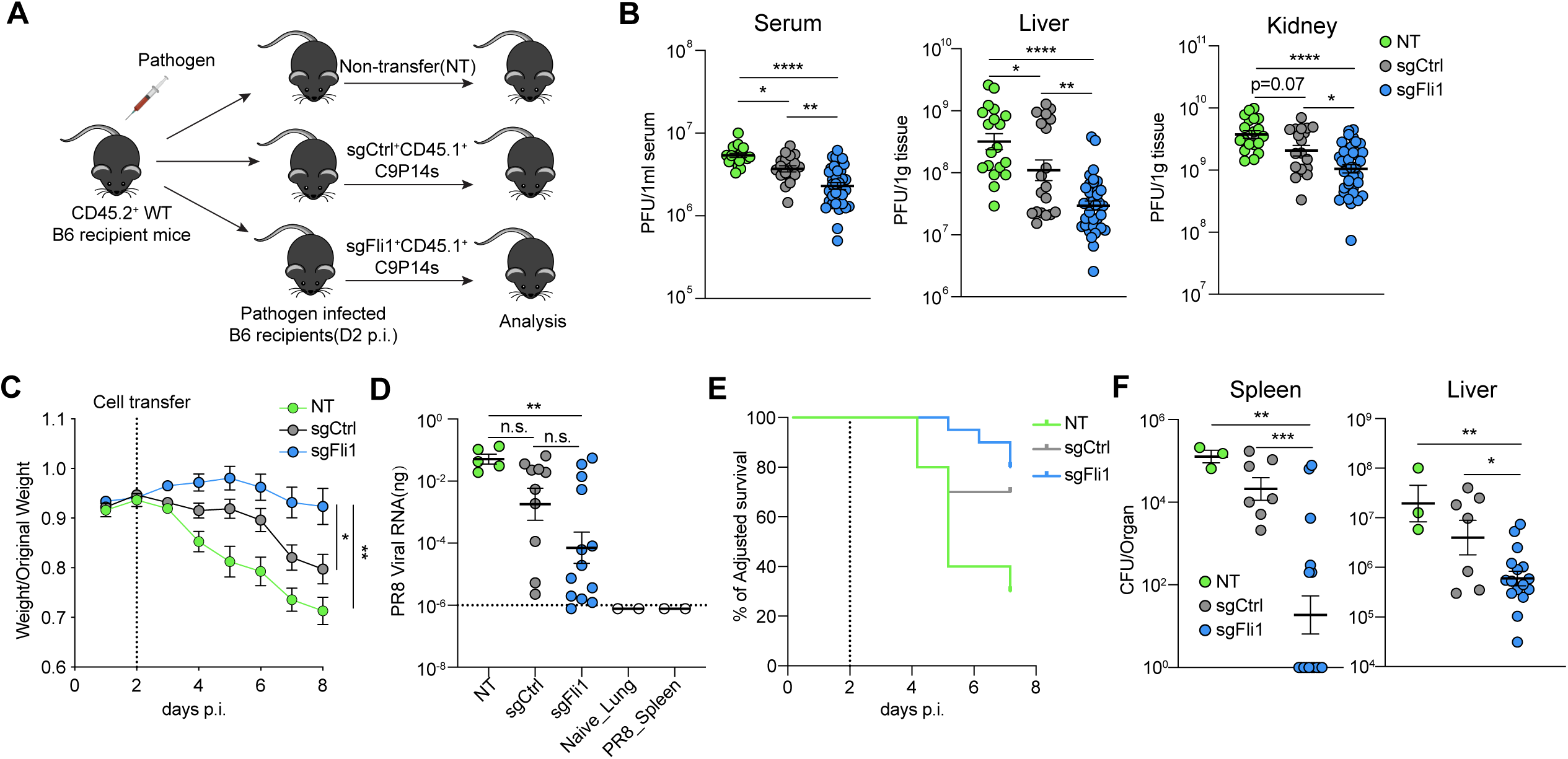
*Fli1*-deficiency in CD8 T cells leads to better protective immunity against infections. A. Experimental design. On D0, CD8 T cells were isolated from CD45.2^+^ C9P14 donor mice and activated; CD45.1^+^ WT recipient mice were infected with LCMV Cl13, lnfluenza virus PR8-GP33 or *Listeria Monocytogenes-*GP33 (LM-GP33). On D1 p.i., activated C9P14 cells were transduced with sgCtrl or sgFli1 RV for 6 hours. On D2 p.i., Cas9^+^sgRNA(VEX)^+^ P14 cells were purified by flow cytometry for sgCtrl or sgFli1 groups, and adoptively transferred into the infected recipient mice. For LCMV-Cl13, 1.5 x 10^5^ VEX^+^ C9P14 cells were adoptively transferred per mouse; for PR8-GP33 and LM-GP33, 1.0 x 10^5^ VEX^+^ C9P14 cells were adoptively transferred per mouse. B. LCMV-Cl13 viral load was measured by plaque assay on D15 p.i. with Cl13 in liver, kidney and serum of non-transferred (NT) mice or mice receiving sgCtrl^+^ or sgFli1^+^ adoptively transferred C9P14 cells. Data pooled from 2 independent experiments. C. Weight curve for PR8-GP33 infected mice from NT, sgCtrl^+^ cell-transferred, or sgFli1^+^ cell-transferred groups. Dashed line represents the time of C9P14 adoptive transfer. D. PR8-GP33 viral RNA load in the lung of NT, sgCtrl^+^ or sgFli1^+^ C9P14 recipient mice. Dashed line indicates the limit of detection. Lung samples from naïve mice and spleen samples from PR8-GP33 infected mice were used as negative controls. E. Adjusted survival curve of LM-GP33 infected mice for NT, sgCtrl^+^ or sgFli1^+^ C9P14 recipient mice. Dashed line represents the time of C9P14 adoptive transfer. F. LM-GP33 bacteria load in spleen and liver of the surviving NT, sgCtrl^+^ or sgFli1^+^ C9P14 recipient mice on D7 p.i. \**P*<0.05, \*\**P*<0.01, \*\*\**P*<0.001, \*\*\*\**P*<0.001 versus control (One-Way Anova analysis). Data are representative of 2 independent experiments (mean±s.e.m.) with at least 3 mice/group.

During LCMV-Cl13 infection, adoptive transfer of sgCtrl^+^ C9P14 cells conferred a moderate degree of viral control compared to the no transfer condition (NT, **Figure 6B**) as expected (Blattman et al., 2009). However, when we compared the two transferred groups, sgFli1^+^ C9P14 provided substantially improved control of viral replication at ∼2 weeks p.i. (**Figure 6B**), demonstrating a phenotypic benefit due to loss of Fli1 even in chronic viral infection where induction of exhaustion is a major barrier to effective protective immunity.

We next evaluated the impact of loss of Fli1 during acutely resolving infections. During influenza-PR8_GP33_ infection, mice receiving sgFli1^+^ C9P14 cells lost less weight than control non-transferred mice or mice receiving sgCtrl^+^ C9P14 cells (**Figure 6C**). This decreased weight loss was associated with better control of viral replication in the lungs of mice receiving sgFli1^+^ but not sgCtrl^+^ C9P14 cells (**Figure 6D**). In this setting, there was variation in the magnitude of the sgFli1^+^ C9P14 expansion in the lungs following PR8_GP33_ infection (**Figure S6A-S6C**). This heterogeneity in the T cell responses was associated with differences in viral control, with some mice nearly eliminating viral RNA by this time point (**Figure 6D**) and recovering from infection-associated weight loss. Therefore, we tested whether there was a relationship between recovery from infection and/or control of viral replication and the magnitude of the C9P14 response. Indeed, the overall magnitude of the C9P14 response was lower in mice that had recovered, consistent with prolonged viral replication and higher antigen load driving increased T cell expansion in mice that had not yet controlled the infection (**Figure S6A-S6B**). Notably, 6 out of 12 of the mice receiving sgFli1^+^ C9P14 cells had controlled disease by this time point, as compared to only 1 out of 11 mice receiving sgCtrl^+^ C9P14 (**Figure 6D, S6B**). In the group of mice still harboring viral RNA in the lungs, sgFli1^+^ C9P14 cells had expanded to substantially higher numbers than sgCtrl^+^ C9P14 cells (**Figure S6B**). A similar difference in T_EFF_ cell expansion was also observed in spleen where sgFli1^+^ C9P14 cells were ∼3 times more numerous than the sgCtrl^+^ group (**Figure S6C**).

Loss of Fli1 conferred a similar advantage following LM_GP33_ infection. Although both sgCtrl^+^ and sgFli1^+^ C9P14 cells improved survival following a high dose LM_GP33_ challenge (**Figure 6E**), sgFli1^+^ C9P14 cells resulted in substantially better control of bacterial replication in the spleen and liver compared to the sgCtrl^+^ C9P14 cells at D7 p.i. (**Figure 6F**). Consistent with the influenza virus setting, this improved protective immunity was associated with greater numerical expansion of sgFli1^+^ C9P14 cells compared to the sgCtrl^+^ group (**Figure S6D**). Thus, deficiency in Fli1 confers a substantial benefit on T_EFF_ cell expansion and protective immunity during chronic LCMV-Cl13 infection, respiratory influenza virus infection, and systemic infection with an intracellular bacterium.

### Loss of Fli1 in CD8 T cells enhances immunity to tumors

We next asked whether the Fli1 deficiency could also mediate increased protective immunity in a cancer setting. Thus, we next employed a subcutaneous B16_GP33_ tumor model. Tumor-bearing mice received equal numbers of sgCtrl^+^ or sgFli1^+^ C9P14 cells on day 5 post tumor inoculation (p.t.) (**Figure 7A**). We used Rag2^-/-^ recipient mice to isolate the effects of sgCtrl^+^ versus sgFli1^+^ C9P14 cells (**Figure 7A**). In this setting sgFli1^+^ C9P14 cells robustly controlled tumor progression compared to non-transferred mice or the sgCtrl^+^ C9P14 group (**Figure 7B**). Furthermore, tumor weight was significantly lower in the sgFli1^+^ C9P14 group compared to either control group at endpoint (**Figure 7C**). Although there was not a clear difference in the number of C9P14 cells/g of tumor, this tumor control was associated with a significant increase in Ly108^-^CD39^+^ donor C9P14 in the sgFli1^+^ group, consistent with a shift towards the more T_EFF_-like population (**Figure 7D-7E**). In the spleen, however, there was a significant increase in sgFli1^+^ C9P14 cell numbers, as well as the proportion of Ly108^-^CD39^+^ cells compared to the sgCtrl^+^ group (**Figure 7F-7G**). We next extended these findings into immune competent mice using B16_GP33_ tumors. We used Cas9^+^ C57BL/6 recipient mice to prevent rejection of the C9P14 donor cells to allow responses to be analyzed for an extended period of time (**Figure S7A**). In this setting, sgFli1^+^ C9P14 cells again conferred substantial benefit on tumor control (**Figure S7B-S7C**) compared to sgCtrl^+^ C9P14 cells. Furthermore, this improved tumor control was associated with an increase in the Ly108^-^CD39^+^ T_EFF_-like population in the tumor, draining lymph node (dLN) and spleen, as well as the increased C9P14 cell numbers in the dLN and spleen (**Figure S7D-S7G**). Thus, genetic deletion of Fli1 conferred a substantial benefit on tumor control indicating a central role for Fli1 in coordinating and restraining protective T_EFF_ responses during tumor invasion and progression. Together, these data show that loss of Fli1 results in improved protective immunity, in the setting of systemic and local, acutely resolving and chronic infections, as well as tumor progression. In these settings enhanced CD8 T cell expansion and a shift in differentiation highlight the role of Fli1 as a key regulator of CD8 T cell differentiation. These data suggest that manipulating Fli1-related pathways could be a viable strategy for therapeutics for infectious disease and cancer.

**Figure 7.**
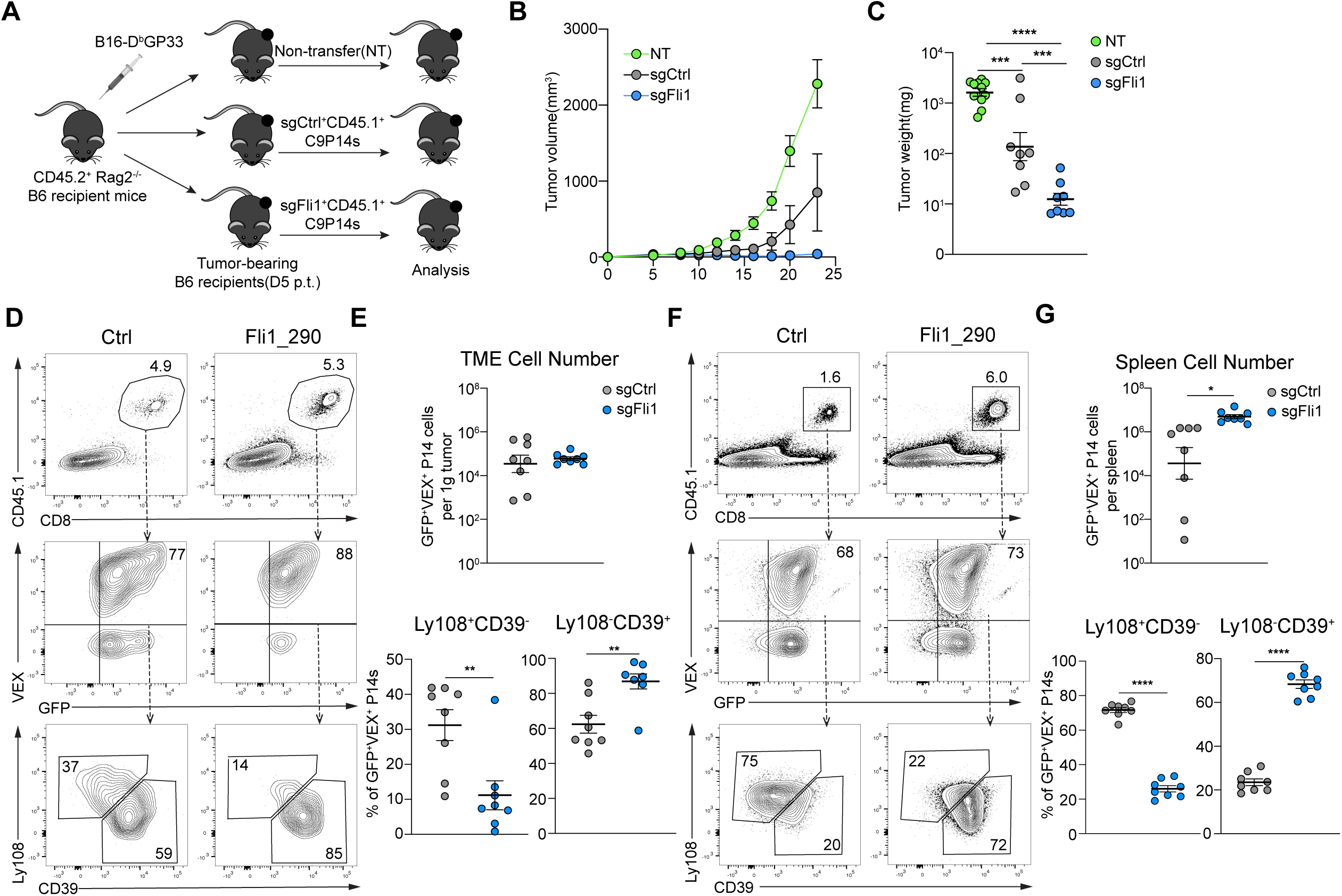
Loss of Fli1 in CD8 T cells results in more potent anti-tumor immunity. A. Experimental design. On D0, CD45.2^+^*Rag2*^-/-^ mice were inoculated with 1 x 10^5^ B16-D^b^gp33 cells. On D3 post tumor inoculation (p.t.), CD8 T cells were isolated from CD45.1^+^ C9P14 mice and activated. On D4 p.t., activated C9P14 cells were transduced with sgCtrl or sgFli1 RVs for 6-8 hours. On D5 p.t., sgRNA (VEX)^+^Cas9 (GFP)^+^P14 cells were purified by flow cytometry from sgCtrl or sgFli1 groups, and 1 x 10^6^ purified VEX^+^ C9P14 cells were adoptively transferred into the tumor-bearing mice. B. Tumor volume curve for mice receiving NT, sgCtrl^+^ or sgFli1^+^ C9P14 cells. C. Tumor weight on D23 p.t. for mice receiving NT, sgCtrl^+^ or sgFli1^+^ C9P14 cells. D-E. Flow cytometry plots (D) and statistical analysis (E) of CD45.1^+^ sgRNA(VEX)^+^Cas9^+^P14 cells and Ly108^-^CD39^+^/Ly108^+^CD39^-^ C9P14 cells from tumor for sgCtrl or sgFli1 groups on D23 p.t. F-G. Flow cytometry plots (F) and statistical analysis (G) of CD45.1^+^ sgRNA^+^Cas9^+^ P14 cells and Ly108^-^CD39^+^/Ly108^+^CD39^-^ C9P14 cells from spleen for sgCtrl or sgFli1 groups at D23 p.t. \**P*<0.05, \*\**P*<0.01, \*\*\**P*<0.001, \*\*\*\**P*<0.001 versus control (two-tailed Student’s *t*-test and One-Way Anova analysis). Data are representative of 2 independent experiments (mean±s.e.m.) with at least 5 mice/group.

## Discussion

Considerable current efforts focus on improving immunotherapy for cancer and chronic infections through a better understanding of the biology of T_EX_ and T_EFF_ differentiation. In this study, we used a focused *in vivo* CRISPR screening approach to specifically interrogate the mechanisms governing T_EX_ versus T_EFF_ differentiation. Among our screening hits, we identified Fli1 as a key TF that safeguards the transcriptional and epigenetic commitment to full T_EFF_ differentiation. Fli1 also negatively regulated CD8 T cell expansion and genetic loss of Fli1 resulted in considerable improvement in antigen-specific CD8 T cell expansion generating more T_EFF_ without numerical loss of progenitor or memory precursor populations. Mechanistically, Fli1 limited epigenetic accessibility to ETS:RUNX sites, preventing Runx3 from fully enabling an effector program. As a result, deleting Fli1 robustly improved protective immunity in multiple models of acute infection, chronic infection and cancer, thus identifying Fli1 as a novel regulator of the T_EFF_ versus T_EX_ differentiation programs and a target for future immunotherapy strategies.

Recent advances in CRISPR-based screening approaches have allowed the dissection of *in vitro* T cell activation (Shang et al., 2018; Shifrut et al., 2018) and *in vivo* responses to infections and tumors (Dong et al., 2019; LaFleur et al., 2019b; Wei et al., 2019; Ye et al., 2019). To better understand the fate commitment of T_EFF_ and T_EX_, we developed OpTICS, an *in vivo* screening system that does not disrupt T cell maturation or the very early signals of T cell activation. This screen revealed known targets essential for positive regulation of the primary steps in T cell activation and differentiation including *Batf, Irf4* (Kurachi et al., 2014; Man et al., 2013) and *Myc* (Lindsten et al., 1988; Wang et al., 2011) as well as TFs that have key roles in developing and controlling the effector program such as members of the NF-κB family and the T-box TF, T-bet. A key goal of this screen was to identify TFs that restrain full T_EFF_ differentiation and prevent optimal protective CD8 T cell responses during chronic infection and cancer. Genes known to promote T cell exhaustion, like *Tox, Nr4a1* and *Nr4a2* (Alfei et al., 2019; Chen et al., 2019a; Khan et al., 2019; Liu et al., 2019; Scott et al., 2019; Seo et al., 2019; Yao et al., 2019) were revealed, as was *Tcf7* due to its the ability to antagonize T cell activation and T_EFF_ differentiation (Chen et al., 2019b). We also, however, identified several novel negative regulators of T_EFF_ differentiation, including *Smad2, Erg* and *Fli1*. Indeed, genetic loss of Fli1 improved protective immunity in multiple settings of acute or chronic infection and cancer. This enhanced protection by *Fli1*-deficient CD8 T cells was robust and consistent across models. Moreover, unlike the effect seen with of the loss T_EX_-driving TF Tox, in which responses cannot be sustained during chronic infection or cancer due to loss of T_EX_ progenitor cells (Khan et al., 2019, Seo et al., 2019), deficiency in Fli1 did not diminish the T_EX_ progenitor population. These observations may be of particular relevance for immunotherapy, where effective immune control is often limited by T cell exhaustion. It remains unknown whether this beneficial effect on tumor immunity is due to increased T cell numbers, de-repressed T_EFF_ differentiation or a combination of these effects. It is also possible that enhanced Runx3 activity fosters more efficient tissue residency, an effect that would be consistent with improved immunity in the influenza virus model. Finally, given the ability to apply CRISPR-mediated genetic manipulations in cellular therapy settings (Stadtmauer et al., 2020; Xu et al., 2019), it is possible to envision improved CAR T cell therapies through targeting Fli1 or related pathways.

Fli1 has a role in hematopoietic stem cell differentiation and co-localizes with other TFs such as Gata1/2 and Runx1 (Tijssen et al., 2011). Here, we find that genetic perturbation of Fli1 significantly increased chromatin accessibility at ETS:RUNX motifs in antigen-specific CD8 T cells responding to viral infection. Moreover, the effect of enforced Runx3 expression is enhanced in the absence of Fli1. These observations suggest that Fli1 prevents accessibility to RUNX binding sites, restricting the activity of the effector-promoting TF Runx3. Moreover, Runx3 can coordinate epigenetic changes at loci encoding other effector-promoting TFs (Wang et al., 2018). Runx1 likely antagonizes Runx3 and vice versa, though Runx3 appears to dominate in settings of T cell activation (Cheng et al., 2008; Cruz-Guilloty et al., 2009). Our data also suggest that Fli1 can cooperate with Runx1 to restrain T_EFF_ differentiation. Together, these data suggest a model where Fli1, in combination with Runx1, prevents efficient genome accessibility or activity of Runx3 and thus restrains a full effector gene program and a positive feed-forward effector promoting activity of Runx3. Thus, genetic deletion of Fli1 de-represses T_EFF_ differentiation, at least partially by creating opportunities for more efficient Runx3 activity.

Recent work has begun to define the transcriptional circuitry that directs fate decisions between terminal T_EFF_, T_MEM_ and T_EX_. Many of these mechanisms that promote one cell fate directly repress the opposing fate. For example, Tox promotes T_EX_ while repressing T_EFF_, TCF-1 promotes T_MEM_ or T_EX_ at the expense of T_EFF_ and Blimp-1, T-bet, Id-2, and others drive T_EFF_ and repress T_MEM_(Kaech and Cui, 2012). The identification of Fli1 as a type of genomic “safeguard” against over-commitment to effector differentiation reveals several novel concepts. First, previous TF circuits that prevent T_EFF_ differentiation to preserve T_MEM_ or T_EX_ differentiation such as TCF-1 or Tox have been essentially binary (Alfei et al., 2019; Chen et al., 2019b; Im et al., 2016; Khan et al., 2019; Utzschneider et al., 2016). Fli1 appears to represent a distinct type of damper on an otherwise robust feed-forward effector transcriptional circuit. By restraining Runx3 node in the effector wiring, Fli1 tempers a central step that not only directly controls expression of key effector genes but also positively reinforces other cooperating effector TFs. However, unlike TCF-1 and Tox, Fli1 does not appear to be required for progenitor biology and the number of both T_EFF_ cells and T_MP_ (in acutely resolved infections) or T_EX_ progenitor cells (in chronic infection) were increased in the absence of Fli1. Thus, by interrupting this “damper” in the circuit, rather than deleting the master switch of T_MP_ or T_EX_ differentiation it may be possible to augment beneficial aspects of short-term protective immunity without compromising long-term immunity. Second, our data reveal a mechanism of competition for epigenetic access between Fli1 and ETS:RUNX family members. These effects may manifest because Fli1 occupies genomic locations that can be bound by ETS:RUNX family TFs that would catalyze chromatin accessibility changes; alternatively, these effects may be due to chromatin changes coordinated by Fli1 itself. It is interesting to note that the EWS-FLI1 fusion recruits the BAF complex to initiate chromatin changes in key regions in cancer cells (Boulay et al., 2017), consistent with the major changes in chromatin accessibility observed in the absence of Fli1 in CD8 T cells here. Thus, the role of Fli1 in CD8 T cells likely involves a chromatin accessibility-based mechanism to restrain ETS:RUNX driven effector biology, though other effects through IRF1/IRF2 may also exist.

The current studies demonstrate a major beneficial effect of loss of Fli1 on protective immunity in multiple settings of acutely resolved or chronic infection and cancer. Although clinical benefit has been observed in some settings due to partial loss of Tox (Khan et al., 2019) or genetic loss of Tox2 with knockdown of Tox (Seo et al., 2019), there may not be substantial benefit for overall protective immunity because complete loss of Tox diverts CD8 T cells entirely towards terminal T_EFF_ differentiation. In contrast, in the absence of Fli1 there was robust and consistent improvement in protective immunity across models. Of particular relevance for immunotherapy, deleting Fli1 improved control of both tumor growth and chronic LCMV infection where the induction of exhaustion typically limits immunity and in the tumor setting. For the former, established B16_gp33_ tumors were effectively controlled for 3 weeks, consistent with the notion that loss of Fli1 provides durable protection in the setting of cancer. It will be interesting to determine in future studies whether this beneficial effect on tumor immunity is due to increased T cell numbers, the de-repression of T_EFF_ differentiation by loss of Fli1, enhanced transcription of specific effector gene programs or a combination of these effects. It is also possible that enhanced Runx3 activity fosters more efficient tissue residency effects, a change that would be consistent with improved immunity in the influenza virus model. Finally, given the ability to apply CRISPR-mediated genetic manipulations in cellular therapy settings (Stadtmauer et al., 2020; Xu et al., 2019) it might be possible to envision clinical benefits with CAR T cells by targeting Fli1 or related pathways.Thus, the OpTICS platform provides a highly robust *in vivo* platform to screen genes involved in regulating of CD8 T cell differentiation as it relates to tumor immunotherapy. This highly focused and optimized platform allows for a 20-100-fold enrichment of sgRNA detection and considerable resolution for gain-of-function screening. In addition to the novel role of Fli1 revealed here, many other potential targets for exploration exist from this screen. Moreover, using OpTICS to extend from this TF focused biology to other areas of cellular biology should provide a robust platform for future discovery.

## Acknowledgement

We thank all members of the Wherry Lab and Shi Lab for helpful discussions. This work was supported by grants from the NIH (AI105343, AI117950, AI082630, AI112521, AI115712, AI108545, CA210944) and Stand Up 2 Cancer to E.J.W.; NIH grant CA234842 to Z.Chen; NIH grant CA009140 to J.R.G.; NIH grant MH109905, HG010480 and Searle Scholar’s Program to A.B.. E.J.W. is supported by the Parker Institute for Cancer Immunotherapy which supports the cancer immunology program at UPenn; S.F.N. is supported by an Australia NH&MRC C.J. Martin Fellowship (1111469) and the Mark Foundation Momentum Fellowship; J.R.G. is supported by Cancer Research Institute-Mark Foundation Fellowship.

## Author Contribution

Z.Chen, J.S. and E.J.W. designed the complete study; Z.Chen performed the mouse experiments, flow cytometry and sorting experiments with the help of Z.Cai, A.E.B.,S.F.N. and E.F.; E.A. and Z.Chen performed sgRNA library construction with the support of J.S., J.R.G., O.K. and S.M.; E.A., E.F., Z.Cao, Z. Wen performed RV-sgRNA vector constructions; Z.Chen performed RV construction, production and transduction with the support from J.E.W., M.K. and N.A.S.; Z.Chen, O.K. and A.E.B. optimized *in vivo* screening conditions; Z.Chen and Y.H. performed sgRNA enrichment analysis with help from J.S. and A.B.; Z.Chen, Y.H. and S.M. performed RNA-seq and analysis; O.K., Z.Chen and S.M. performed ATAC-seq and analysis; Z.Z., H.H and Z.Chen performed CUT&RUN and analysis with the support of S.L.B.; Z.Chen performed LM infection; M-A. A. and Z.Chen performed influenza infection and detection assays; Z.Chen, K.N. and V.E. performed plaque assays of LCMV-Cl13; Z.Chen and S.F.N. performed B16 tumor experiments; A.G., A.E.B. and J.E.W. made key editorial contributions to the manuscript; Z.Chen, J.S. and E.J.W. wrote the manuscript.

## STAR⋆Methods

### KEY RESOURCES TABLE

| REAGENT or RESOURCE | SOURCE | IDENTIFIER |
| --- | --- | --- |
| **Mouse strains** |  |  |
| C57BL/6 | Charles River | N/A |
| CD45.1<sup>+</sup> C57BL/6 | Charles River | N/A |
| CD4<sup>CRE</sup> | The Jackson Lab | Stock No. 022071 |
| TCRα<sup>-/-</sup>; P14 TCRVα2Vβ8 | The Jackson Lab | Stock No. 37394-JAX |
| Rag2<sup>-/-</sup> C57BL/6 | The Jackson Lab | Stock No: 008449 |
| LSL-Cas9-GFP | The Jackson Lab | Stock No: 026175 |
| Constitutive-Cas9-GFP | The Jackson Lab | Stock No: 026179 |
| **Flow cytometry reagents** |  |  |
| Live/Dead Aqua Dye | Thermofisher | Cat# L34957 |
| Live/Dead Zombie NIR Dye | BioLegend | Cat# 423106 |
| Anti-Mouse KLRG1(2F1) | BD Biosciences | Cat# 561619, RRID:AB_10898017 |
| Anti-Mouse CD127(A7R34) | BioLegend | Cat# 135016, RRID:AB_1937261 |
| Anti-Mouse CD8(53-6.7) | BioLegend | Cat# 100742, RRID:AB_2563056 |
| Anti-Mouse CD44(IM7) | BioLegend | Cat# 103059, RRID:AB_2571953 |
| Anti-Mouse CD45.1(A20) | BioLegend | Cat# 110724, RRID:AB_493733;<br>Cat# 110716, RRID:AB_313505 |
| Anti-Mouse CD45.2(104) | BioLegend | Cat# 109828, RRID:AB_893350;<br>Cat# 109823, RRID:AB_830788 |
| Anti-Mouse Ly108(330-AJ) | BioLegend | Cat# 134608, RRID:AB_2188093;<br>Cat# 134605, RRID:AB_1659258 |
| Anti-Mouse Tim-3(RMT3-23) | BioLegend | Cat# 119721, RRID:AB_2616907 |
| Anti-Mouse CD39(24DMS1) | Thermofisher | Cat# 46-0391-80, RRID:AB_10717513 |
| Anti-Mouse PD-1(RMP1-30) | BioLegend | Cat# 109109, RRID:AB_572016 |
| Anti-Mouse CD28 | BioLegend | Cat# 644808, RRID:AB_1595479 |
| Anti-Mouse CX3CR1 | Thermofisher | Cat# 50-4875-80, RRID:AB_2574226 |
| Anti-Mouse CXCR3 | BioLegend | Cat# 652420, RRID:AB_2564285 |
| Anti-Mouse TCF-1(S33-966) | BD Biosciences | Cat# 564217, RRID:AB_2687845 |
| Anti-Mouse Gzmb (GB11) | Thermofisher | Cat# GRB17, RRID:AB_2536540 |
| LCMV D<sup>b</sup>GP33 tetramer | NIH | Conjugated in house |
| Foxp3 Transcription Factor<br>Staining Buffer Kit | Thermofisher | Cat# A25866A |
| **Experimental Models** |  |  |
| LCMV Clone13 (CI13) | Rafi Ahmed | Grew up in house |
| LCMV Armstrong (Arm) | Rafi Ahmed | Grew up in house |
| *Listeria Monocytogenes*-D<sup>b</sup>GP33 | Hao Shen | Grew up in house |
| *Influenza-PR8*-D<sup>b</sup>GP33 | Richard J. Webby | Grew up at St. Jude Children's Hospital |
| B16-D<sup>b</sup>GP33 | (Prévost-Blondel et al.,<br>1998a) | Grew up in house |
| **In vitro culture and retroviral transduction reagents** |  |  |
| Recombinant human IL-2 | NIH | N/A |
| Anti-Mouse CD3(145-2C11) | BioLegend | Cat# 100302, RRID:AB_312667 |
| Anti-Mouse CD28(37.51) | Thermofisher | Cat# 16-0281-82, RRID:AB_468921 |
| EasySep™ Mouse CD8+ T Cell<br>Isolation Kit | STEMCELL Technologies | Cat# 19853 |
| RPMI-1640 medium | Corning/Mediatech | Cat# 10-040-CV |
| DMEM medium | Corning/Mediatech | Cat# 10-017-CV |
| HI Fetal Bovine Serum | Thermofisher | Cat# 26170-043 |
| HEPES | Thermofisher | Cat# 15630080 |
| Non-Essential Amino Acids | Thermofisher | Cat# 11140050 |
| Penicillin-Streptomycin | Thermofisher | Cat# 15140122 |
| β-mercaptoethanol | Sigma-Aldrich | Cat# M6250-500ML |
| Opti-MEM | Thermofisher | Cat# 31985088 |
| Polybrene | Sigma-Aldrich | Cat# TR-1003-G |
| Lipofectamine™ 3000 Transfection<br>Reagent | Thermofisher | Cat# L3000001 |
| **Molecular constructs** |  |  |
| Runx1 overexpression vector | Nancy A. Speck; Addgene | N/A; Addgene Cat#80157 |
| Runx3 overexpression vector | In this paper | N/A |
| Fli1 overexpression vector | In this paper | N/A |
| Empty-VEX retroviral vector | (Kurachi et al. 2017) | N/A |
| Empty-mCherry retroviral vector | (Kurachi et al. 2017) | N/A |
| pSL21-VEX | In this paper | N/A |
| pSL21-mCherry | In this paper | N/A |
| LRG2.1 | Addgene | Cat#108098 |
| MSCV Retroviral Expression System | Takara Bio. | Cat#634401 |
| BbsI | NEB | Cat#R0539L |
| Esp3I(BsmBI) | Thermofisher | Cat#ER0451 |
| Phusion Flash High Fidelity PCR Master Mix with HF buffer | Thermofisher | Cat#F531L |
| Gibson Assembly Master Mix | NEB | Cat#E2611L |
| T4 DNA Polymerase | NEB | Cat#M0203L |
| DNA Polymerase I, Large (Klenow) Fragment | NEB | Cat#M0210L |
| T4 polynucleotide kinase | NEB | Cat#M0201L |
| Klenow Fragment | NEB | Cat#M0212L |
| PrimeSTAR HS DNA Polymerase | Takara Bio. | Cat#R040A |
| Quick T4 DNA Ligase | NEB | Cat#M2200 |
| **Antibodies for biochemical experiments** |  |  |
| Anti-Mouse Fli1 | Abcam | Cat# ab15289 |
| Guinea Pig anti-Rabbit IgG (Heavy & Light Chain) Antibody | Antibodies-online | Cat# ABIN101961 |
| Anti-Mouse GAPDH | Cell Signaling Technology | Cat# 14C10 |
| IRDye 800 CW Goat anti-Rabbit Antibody | Licor | #92632211 |
| **RNA-Sequencing Processing Reagents** |  |  |
| RNeasy Micro Kit | Qiagen | Cat# 74004 |
| SMART-Seq® v4 Ultra® Low Input RNA Kit for Sequencing (24 rxn) | Takara Bio | Cat# 634889 |
| Nextera XT DNA Library Preparation Kit | Illumina | Cat# FC-131-1024 |
| Agencourt AMPure XP | Beckman | Cat# A63880 |
| HSD5000 ScreenTape | Agilent | Cat# 5067-5592 |
| HSD1000 ScreenTape | Agilent | Cat# 5067-5584 |
| **ATAC-Sequencing Processing Reagents** |  |  |
| IGEPAL-CA-630 | Sigma-Aldrich | Cat# I8896 |
| Tn5 Transposes | Illumina | Cat# FC-121-1030 |
| MinElute PCR Purification Kit | Qiagen | Cat# 28004 |
| NEBNext High-Fidelity 2x PCR Master Mix | New England Labs | Cat# M0541 |
| **CUT&RUN Processing Reagents** |  |  |
| Digitonin | Millipore | Cat# 300410-1GM |
| BioMag Plus Concanavalin A (10mL) | Bangs laboratories | Cat# BP531 |
| Complete, EDTA-free Protease Inhibitor Cocktail | Sigma | Cat# 4693132001 |
| Glycogen | Thermo | Cat# R0561 |
| Proteinase K | Denville Scientific | Cat# CB3210-5 |
| RNase A | Thermo | Cat# EN0531 |
| Spermidine | Sigma | Cat# 85558-1G |
| NEB Next Ultra II DNA library prep kit | NEB | Cat# E7645L |
| **Illumina Sequencing Reagents** |  |  |
| NEBNext Library Quant Kit for Illumina | NEB | Cat# E7630L |
| NextSeq 500/550 High Output Kit (75 cycles) v2.5 kit | Illumina | Cat# 20024906 |
| NextSeq 500/550 High Output Kit (150 cycles) v2.5 kit | Illumina | Cat# 20024907 |
| **Software and Algorithms** |  |  |
| FlowJo v10.4.2 | TreeStar | https://www.flowjo.com/solutions/flowjo/downloads |
| Prism Version 8 | GraphPad Software | https://www.graphpad.com/scientific-software/prism/ |
| IGV v2.4.16 | The Broad Institute | https://software.broadinstitute.org/software/igv/download |
| Deseq2 | (Love et al., 2014) | http://www.bioconductor.org/packages/release/bioc/html/DESeq2.html |
| Great (v3.0) | (McLean et al., 2010) | http://great.stanford.edu/public/html/ |
| GSEA (v3.0) | The Broad Institute | http://software.broadinstitute.org/gsea/index.jsp |
| ClusterProfiler | (Yu et al., 2012) | https://guangchuangyu.github.io/2016/01/go-analysis-using-clusterprofiler/ |
| Homer (v4.6) | (Heinz et al., 2010) | http://homer.ucsd.edu/homer/download.html |
| Bowtie2 v2.3.4.1 | (Langmead and Salzberg, 2012) | http://bowtie-bio.sourceforge.net/bowtie2/index.shtml |
| Picard v1.96 | The Broad Institute | https://broadinstitute.github.io/picard/index.html |
| Samtools v1.1 | (Li et al., 2009) | http://samtools.sourceforge.net |
| Bedtools v2.28.0 | (Quinlan and Hall, 2010) | https://bedtools.readthedocs.io/en/latest/# |
| bedGraphToBigWig (UCSC) | UCSC Genome | http://hgdownload.soe.ucsc.edu/admin/exe/linux.x86_64/ |
| MACS v2.1 | (Zhang et al., 2008) | https://github.com/taoliu/MACS |
| HOMER v4 | (Heinz et al., 2010) | http://homer.ucsd.edu/homer/ |
| ChIPpeakAnno | (Zhu et al., 2010) | https://bioconductor.org/packages/release/bioc/html/ChIPpeakAnno.html |
| ComplexHeatmap | (Gu et al., 2016) | https://bioconductor.org/packages/release/bioc/html/ComplexHeatmap.html |
| **Datasets** |  |  |
| Blacklist | ENCODE | https://sites.google.com/site/anshulkundaje/projects/blacklists |
| T<sub>EFF</sub> gene signature | (Bengsch et al., 2018) | N/A |
| T<sub>EX</sub> precursor gene signature | (Chen et al., 2019b) | GSE131535 |
| sgCtrl vs sgFli1 RNA sequencing at D8 CI13 p.i. | In this paper | GSE149838 |
| sgCtrl vs sgFli1 ATAC sequencing at D9 CI13 p.i. | In this paper | GSE149836 |
| IgG vs Fli1-ab(ab15289)<br>CUT&RUN Sequencing at D8 CI13 p.i. | In this paper | GSE149837 |

## Experimental Procedures

### Mice

CD4^CRE^, LSL-Cas9-GFP and Constitutive-Cas9-GFP mice were purchased from Jackson Laboratory. LSL-Cas9-GFP mice were bred to CD4^CRE^ mice and TCR transgenic P14 C57BL/6 mice (TCR specific for LCMV D^b^GP_33–41_) and back crossed for more than 6 generations before use. Constitutive-Cas9-GFP mice were bred to TCR transgenic P14 C57BL/6 mice. Constitutive-Cas9-GFP mice for recipient use were bred in house. 6-8 week-old C57BL/6 Ly5.2CR (CD45.1) or C57BL/6 (CD45.2) mice were purchased from NCI. 5-7 week-old Rag2^-/-^ C57BL/6 mice were purchased from Jackson Laboratory. Both male and female mice were used. All mice were used in accordance with Institutional Animal Care and Use Committee guidelines for the University of Pennsylvania.

### Experimental models

#### LCMV Infection

Mice were infected intraperitoneally (i.p.) with 2 × 10^5^ plaque-forming units (PFU) LCMV Armstrong or intravenously (i.v.) with 4 × 10^6^ PFU LCMV Cl13. Plaque assay for LCMV-Cl13 to detect viral load was processed as previously described(Pauken et al., 2016).

#### *Listeria Monocytogenes* (*LM*) infection

*LM* expressing D^b^GP33(*LM*-gp33) concentration was measured by optical density (OD) after overnight culture in brain heart infusion (BHI) media (1 OD refers to 8 × 10^8^ *LM*-gp33). Each recipient mouse was infected intravenously (i.v.) with 1 × 10^5^ CFU *LM*-gp33. Adjusted survival was based on mice remaining above the mandatory Institutional Animal Care and Use Committee (IACUC) euthanasia cut off of 30% weight loss. *LM*-gp33 colony formation per unit calculation for bacteria load was calculated as previously described(Chen et al., 2017).

#### Influenza *PR8* infection

Mice were infected intranasally (i.n.) with *PR8* strain expressing D^b^GP33 (*PR8*-gp33) at a dose of 3.0 LD50. Mice were anesthetized before i.n. infection. PR8 viral qPCR detection for viral RNA amount was calculated as previously described(Laidlaw et al., 2013).

#### Tumor transfer

B16F10 melanoma cells expressing D^b^GP33 (B16F10-gp33,(Prévost-Blondel et al., 1998b)) were maintained at 37 °C in DMEM medium supplemented with 10% FBS, penicillin, streptomycin and L-glutamine. Tumor cells were injected subcutaneously into the flanks of Rag2^-/-^mice at 1 x 10^5^ cells/recipient and of Cas9^+^ B6 mice at 2 x 10^5^ cells/recipient. Activated sgRNA^+^ C9P14 cells are sorted and transferred into recipient mice at a dose of 1 x 10^6^ cells/recipient (for Rag2^-/-^) or 3 x 10^6^ cells/recipient (for Cas9^+^). Tumor size was measured using digital calipers every 2-3 days after inoculation.

#### Vector construction and sgRNA cloning

In this study, SpCas9 sgRNA was expressed using pSL21-VEX or pSL21-mCherry (U6-sgRNA-EFS-VEX or U6-sgRNA-EFS-mCherry, will be available through Addgene). To generate the pSL21-VEX or pSL21-mCherry, U6-sgRNA expression cassette was PCR cloned from LRG2.1 into a retroviral vector MSCV-Neo, followed by swapping the Neo selection marker with a VEX or mCherry fluorescence reporter. sgRNAs were cloned by annealing two DNA oligos and T4 DNA ligation into a Bbs1-digested pSL21-VEX or pSL21-mCherry vector. To improve U6 promoter transcription efficiency, an additional 5’ G nucleotide was added to all sgRNA oligo designs that did not already start with a 5’ G. Runx1 and Runx3 constructs are built on the MIGR or MSCV-mCherry constructs, empty MIGR or MSCV-mCherry are used as the controls for these vectors.

#### Cell culture and in vitro stimulation

CD8 T cells were purified from spleens by negative selection using EasySep Mouse CD8^+^ T Cell Isolation Kit (STEMCELL Technologies) according to manufacturer’s instructions. Cells were stimulated with 100 U/mL recombinant human IL-2, 1 µg/mL anti-mouse CD3ε, and 5 µg/mL anti-mouse CD28 in RPMI-1640 medium with 10% fetal bovine serum (FBS), 10 mM HEPES, 100 µM non-essential amino acids (NEAA), 50 U/mL penicillin, 50 µg/mL streptomycin, and 50 µM β-mercaptoethanol.

#### Retroviral vector (RV) experiments

RVs were produced in 293T cells with MSCV and pCL-Eco plasmids using Lipofectamine 3000. RV transduction was performed as described (Kurachi et al., 2017). Briefly, CD8^+^ T cells were purified from spleens of P14 mice using EasySep™ Mouse CD8^+^ T Cell Isolation Kit. After 18-24 hrs of *in vitro* stimulation, P14 cells were transduced with RV in the presence of polybrene (0.5 µg/ml) during spin infection (2,000 g for 60 min at 32°C) following incubation at 37°C for 6 hrs for single RV and sgRNA library, or 12 hrs for double RV. RV-transduced P14 cells were adoptively transferred into recipient mice that were infected 24-48 hrs prior to transfer.

#### Flow cytometry and sorting

For mouse experiments, tissues were processed, single cell suspensions obtained, and cells were stained as described (Wherry et al., 2003). Mouse cells were stained with LIVE/DEAD cell stain (Invitrogen) and antibodies targeting surface or intracellular proteins. Flow cytometry was performed with an LSRII. Cell sorting experiments were performed with a BD-Aria sorter, with 70-micron nozzle and a 4°C circulating cool-down system for sequencing, western and TIDE assays.

For sorting RV^+^ cells optimized sorting in the transfer experiments, the BD Aria Sorter was set at 37°C and 100-micron nozzle, with a flow rate lower than 3.0. 3 × 10^6^ Cells are concentrated in 300ul 10% complete RPMI with 100 U/mL recombinant human IL-2 during sorting. 37°C pre-warmed collection tubes with 10% complete RPMI (100 U/ml IL-2) are used. Sorted cells are washed by 37°C warm pure RPMI before transferring into recipients.

#### TIDE Assay

At least 1 x 10^4^ Cas9^+^sgRNA^+^ T cell pellets were frozen down. Genomic DNA was isolated from these samples using QIAmp DNA Mini Kit. A TIDE PCR, using 2x Phusion Flash High-Fidelity PCR Master Mix and primers designed around the genome region of the sgRNA target part was run for each sample to extract the guide region from the genome DNA; the resulting products were then gel verified, PCR purified, and sent for Sanger sequencing.

#### Western Blot

2 x 10^5^ T cells were sorted using FACS machine and the pellets were frozen down. Protein was from these samples was extracted and denatured by boiling at 95°C in 2X working loading sample buffer (1M Tris-HCl, 10% SDS, Glycerol, 10% Bromophenol blue). Lysate was run on a 10% SDS-PAGE gel and then transferred to a nitrocellulose membrane. Primary Fli1 (1:200) and GAPDH (1:1000) antibodies was staining over night, followed 1:5000 secondary antibody staining on the next day.

#### OpTICS screening

- **sgRNA candidate selection** 271 TFs that met the following criteria were selected 1) Among the top 50 differentially expressed across (Doering et al., 2012) and (Philip et al., 2017), 2) Among the top 10 differentially open TF motifs across Naïve, D8 Arm and D8 Cl13 in the previous described(Sen et al., 2016), 3) Involved in the top immune-regulatory families, such as IRF and STAT proteins. 120 TFs were manually chosen to be included in the TF library.
- **Library construction** 4-5 sgRNA were designed against individual DNA binding domains or other functional domains of each TF based on the domain sequence information retrieved from NCBI Conserved Domains Database. All of the sgRNA oligos, including positive and negative control sgRNAs, were synthesized by Integrated DNA Technologies (IDT) and pooled in equal molarity. The pooled sgRNA oligos were then amplified by PCR and cloned into BsmBI-digested SL21 vector using Gibson Assembly Kit. To verify the identity and relative representation of sgRNAs in the pooled plasmids, a deep-sequencing analysis was performed on a MiSeq instrument. We confirmed that 100% of the designed sgRNAs were cloned in the SL21 vector and the abundance of >95% of individual sgRNA constructs was within 5-fold of the mean (data not shown).
- **Mouse Experimental Workflow** On day 0, C9P14 cells were isolated from the spleens and lymph nodes of CD45.2^+^ C9P14 mice and processed to standard T cell activation protocol using anti-CD3/CD28 and IL-2; on the same day, naïve CD45.1^+^ recipient mice were infected by LCMV. On D1 p.i., activated C9P14 cells were transduced by RV-sgRNA library and incubated for 6 hours before washing out the RV supernatant. 18-24 hours later, the transduced sgRNA^+^Cas9^+^ cells were sorted. Then, 10% of the sgRNA^+^Cas9^+^ T cells were frozen down as a D2 baseline (T0 time point) control prior to any selection, while 90% of the cells are transfer to the infected recipients (maximum 1×10^5^ cells/recipient). On the T1 time point (D8 in the graph), sgRNA^+^Cas9^+^ CD45.2^+^ T cells were sorted out from multiple organs of the recipients.
- **Isolated library construction and MiSeq processing** To quantify the sgRNA abundance of reference and end time points, the sgRNA cassette was PCR amplified from genomic DNA using high-fidelity polymerase. The PCR product was end-repaired by T4 DNA polymerase, DNA Polymerase I, Large (Klenow) Fragment, and T4 polynucleotide kinase. Next, a 3’ A-overhang was then added to the ends of blunted DNA fragments with Klenow Fragment (3’-5’ exo-). The DNA fragments were ligated to diversity-increased custom barcodes with Quick ligation kit. Illumina paired-end sequencing adaptors were attached to the barcoded ligated products through PCR reaction with high-fidelity polymerase. The final product was quantified by Bioanalyzer Agilent DNA 1000 and pooled together in equal molar ratio and pair-end sequenced by using MiSeq (Illumina) with MiSeq Reagent Kit V3 150-cycle (Illumina).
- **Data processing** The sequencing data was de-multiplexed and trimmed to contain only the sgRNA sequence cassettes. The read count of each individual sgRNA was calculated with no mismatches and compared to the sequence of reference sgRNA as described previously(Shi et al., 2015). Data of each sample were normalized to the same read account. Waterfall plots (Figure 1B): For each gene, mean of the log2 fold changes from multiple sgRNA was computed. Heatmap (Figure 1C): For each gene, mean of the log10 fold changes from multiple sgRNA was computed. In the matrix of genes by conditions, quantile normalization was performed across conditions such that each condition has the same distribution of values. Genes were ordered by the mean value of each row. Histogram (Figure 1D): For each condition, the background (grey bars and the histogram) was plotted for sgRNAs of all the genes. The 5% and 95% intervals were extracted by using the 5th percentile and 95th percentile of background values. The red bars show the log folds change for sgRNAs for one gene (or the control)

#### RNA-Sequencing

- **Experiment workflow** At D8 p.i. with Cl13, CD8 T cells were isolated from spleens of infected recipients. VEX^+^GFP^+^ cells are sorted using FACS with >95% purity. RNA were isolated using the QIAGEN RNeasy Micro Kit with 2 x 10^4^ cell per sample. cDNA libraries were generated using SMARTSeq V4 Ultra Low kit. Libraries were quantified by qPCR using a KAPA Library Quant Kit (KAPA Biosystems). Normalized libraries were pooled, diluted to 1.8pg/ml loaded onto a TG NextSeq 500/550 Mid Output Kit v2 (150 cycles, 130M reads, Illumina) and paired-end sequencing was performed on a NextSeq 550 (Illumina). The estimated read depth per sample is 15M reads.
- **Data processing** Raw FASTQ files from RNAseq paired-end sequencing were aligned to the GRCm38/mm10 reference genome using Kallisto (https://pachterlab.github.io/kallisto/). Sequencing reads were read in for 19357 genes and 8 samples. Genes with zero read count in more than three conditions were filtered out. 13628 genes remained after this step. Then, differential expression analysis was run using DESeq 2 package. The expression of 1440 genes were found to significantly differ between the two conditions at a BH corrected P-value < 0.05. GO enrichment analysis was performed using ClusterProfiler. The top 20 most enriched pathways are shown in the plot. GSEA was performed using the Broad Institute software (https://www.broadinstitute.org/gsea/index.jsp). Enrichment scores were calculated by comparing sgCtrl to sgFli1 groups. T_EX_ precursor gene signature was from(Chen et al., 2019b). T_EFF_ gene signature was from(Bengsch et al., 2018).

#### ATAC-Sequencing

- **Experimental Workflow** ATACseq sample preparation was performed as described with minor modifications (Buenrostro et al., 2013). VEX^+^GFP^+^ cells were sorted using FACS with >95% purity. Sorted cells (2.5 x 10^4^) were washed twice in cold PBS and resuspended in 50μl of cold lysis buffer (10nM Tris-HCl, pH 7.4, 10mM NaCl, 3mM MgCl2, 0.1% Tween). Lysates were centrifuge (750xg, 10min, 4°C) and nuclei were resuspended in 50μl of transposition reaction mix (TD buffer [25μl], Tn5 Transposase [2.5μl], nuclease-free water [22.5μl]; (Illumina)) and incubated for 30min at 37°C. Transposed DNA fragments were purified using a Qiagen Reaction MiniElute Kit, barcoded with NEXTERA dual indexes (Illumina) and amplified by PCR for 11 cycles using NEBNext High Fidelity 2x PCR Master Mix (New England Biolabs). PCR products were purified using a PCR Purification Kit (Qiagen) and amplified fragment sizes were verified on a 2200 TapeStation (Agilent Technologies) using High Sensitivity D1000 ScreenTapes (Agilent Technologies). Libraries were quantified by qPCR using a KAPA Library Quant Kit (KAPA Biosystems). Normalized libraries were pooled, diluted to 1.8pg/ml loaded onto a TG NextSeq 500/550 Mid Output Kit v2 (150 cycles,130M reads, Illumina) and paired-end sequencing was performed on a NextSeq 550 (Illumina). Estimation read depth per sample is 10M reads.
- **Data processing** Raw ATACseq FASTQ files from paired-end sequencing were processed using the script available at the following repository (https://github.com/wherrylab/jogiles_ATAC). Samples were aligned to the GRCm38/mm10 reference genome using Bowtie2. We used samtools to remove unmapped, unpaired, mitochondrial reads. ENCODE blacklist regions were also removed (https://sites.google.com/site/anshulkundaje/projects/blacklists). PCR duplicates were removed using Picard. Peak calling was performed using MACS v2 (FDR q-value 0.01). For each experiment, we combined peaks of all samples to create a union peak list and merged overlapping peaks with BedTools *merge*. The number of reads in each peak was determined using BedTools *coverage.* Differentially accessible regions were identified following DESeq2 normalization using an FDR cut-off < 0.05 unless otherwise indicated. Motif enrichment was calculated using HOMER (default parameters) on peaks differentially accessible across sgCtrl group and sgFli1 group. Transcription binding site prediction analysis was performed using known motif discovery strategy.

#### CUT&RUN

- **Experimental Workflow** CUT&RUN experiments were performed as previously described(Skene et al., 2018) with modifications. Briefly, 2×10^5^ sorted cells were washed twice with 1 ml of cold wash buffer (20 mM HEPES-NaOH pH 7.5, 150 mM NaCl, 0.5 mM Spermidine, and protease inhibitor cocktails from Sigma) in 1.5ml tubes. Cells were then resuspended in 1 ml of cold wash buffer and incubated with 10 μl of BioMagPlus Concanavalin A (Bangs laboratories) by rotating at 4°C for 25 min to allow the cells to bind. Tubes were placed on a magnetic stand and liquid was removed after the solution turned clear. Primary antibody in 250 μl of cold antibody buffer (20 mM HEPES-NaOH pH 7.5, 150 mM NaCl, 0.5 mM Spermidine, 2 mM EDTA, 0.1% digitonin, and protease inhibitor cocktails from Sigma) was added to the tubes and rotated at 4°C overnight. The next day, after washing cells once with 1 ml of cold wash buffer, protein A-MNase (pA-MN) in 250 μl of cold digitonin buffer (20 mM HEPES-NaOH pH 7.5, 150 mM NaCl, 0.5 mM Spermidine, 0.1% digitonin, and protease inhibitor cocktails from Sigma) was added to the tube and rotate at 4°C for 1 h. To wash away unbound pA-MN, cells were washed twice with 1 ml of cold digitonin buffer, and then resuspended in 150 μl of cold digitonin buffer. The tubes were placed on a pre-cooled metal block. To initiate pA-MN digestion, 3 μl of 0.1 M CaCl_2_ was mixed with cells in 150 μl cold digitonin buffer by gently flicking the tubes 10 times. Tubes were immediately placed back in the metal block. After 30 min incubation, the digestion was stopped by adding 150 μl of 2x stop buffer (340 mM NaCl, 20 mM EDTA, 4 mM EGTA, 0.02% Digitonin, 50 μg/ml RNase A, 50 μg/ml Glycogen, and 4 pg/ml yeast heterologous spike-in DNA). Target chromatin was released by incubating the tubes on a heat block at 37°C for 10 min. Supernatant was spun at 16,000 g for 5 min at 4°C and transferred to a new tube. Chromatin was incubated with 3 μl of 10% SDS and 2.5 μl of 20 mg/ml proteinase K at 70°C for 10 min, followed by phenol:chloroform:isoamyl alcohol extraction. Upper phase containing DNA was mixed with 20 μg of glycogen and incubated with 750 μl of cold 100% ethanol at -20°C overnight. DNA was precipitated by centrifugation at 20,000 g for 30 min at 4°C. DNA pellets was washed once by cold 100% ethanol, air-dried, and stored at -20°C for library preparation. Protein A-MNase (batch 6, use at 1:200) and yeast heterologous spike-in DNA were kindly provided by Dr. Steve Henikoff. The antibodies used were: Fli1, ab15289, used at 1:50 (abcam) and guinea pig anti-rabbit IgG, used at 1:100, ABIN101961 (antibodies-online).
CUT&RUN DNA library was prepared as previously descried(Liu et al., 2018) with slight modifications. Briefly, all DNA precipitated from pA-MN digestion was used for library preparation using NEBNext Ultra II DNA Library Prep Kit (NEB). The adaptor was diluted to 1:25 for adaptor ligation. DNA was barcoded and amplified for 14 PCR cycle, and DNA library was cleaned up by AMPure XP beads(Liu et al., 2018). The library quality was checked with Qubit and bioanalyzer, and the quantity of the library was determined by qPCR using NEBNext Library Quant Kit for Illumina (NEB) according to manufacture’s instruction. Eighteen barcoded libraries were pooled at equal molarity and sequenced in the NextSeq 550 platform with NextSeq 500/550 High Output Kit (75 cycles) v2.5 kit. Paired-end sequencing was carried out (42:6:0:42).
- **Data processing** Paired-end reads were aligned to mm10 reference genome using Bowtie2 v2.3.4.1 with options suggested by Henikoff (Skene et al., 2018). Picard tools v1.96 was used to remove presumed PCR duplicates using the MarkDuplicates command. Bam files containing uniquely mapped reads were created using Samtools v1.1. For downstream analysis, biological replicates (3 per condition) were merged at this step. Bedtools v2.28.0 was used to generate fragment BED files with size 40bp-500bp. Blacklist regions, random chromosomes, and mitochondria were removed. Filtered BED files were used for downstream analysis. Read per million (RPM) normalized bigwig files were created using bedGraphToBigWig (UCSC) and were used to visualize binding signals. Peaks were called using MACS v2.1 using the broadPeak setting with p-value cutoff of 1e-8, -f BEDPE and IgG as controls. Genes proximal to peaks were annotated against the mm10 genome using annotatePeaks.pl from HOMER v4. Fli1 binding motifs were identified using findMotifsGenome.pl from HOMER v4. Venn diagram of comparison with ATAC-Seq peaks was plotted using Bioconductor package ChIPpeakAnno. Heatmap was generated using Bioconductor package ComplexHeatmap.

#### Statistical analysis

Statistical significance was calculated with unpaired two-tailed student’s t-test or one-way ANOVA with Tukey’s multiple comparisons test by Prism 7 (GraphPad Software). P values are reported in the figure legends.

## Supplementary Figure Legends

**Figure S1 High-efficiency gene editing using retroviral-transduced sgRNA in Cas9**^**+**^ **antigen specific CD8 T cells.**

A. Optimizing sgRNA backbone compared to the original sgRNA.

B. Experimental design for *in vivo* gene editing test. At Day 0 (D0), CD8 T cells were isolated from CD45.1^+^ LSL-Cas9^+^CD4^CRE+^P14 (C9P14) donor mice and activated with anti-CD3, anti-CD28 and IL-2; CD45.2^+^ WT recipient mice were infected with LCMV-Cl13. At D1 p.i., activated C9P14 cells were transduced with either Ctrl-sgRNA(sgCtrl) or *Pdcd1*-sgRNA (sgPdcd1). 6 hours after transduction, 5 × 10^4^ activated donor cells were adoptively transferred into infected recipient mice. C9P14 cells were then isolated, at the indicated times, from different organs of recipient mice for analysis.

C. D2 *in vitro* transduction efficiency of activated C9P14 cells with sgRNA vector (mCherry). Gates are set based on the non-transduced control.

D. Flow cytometry plots of Cas9(GFP)^+^sgRNA(mCherry)^+^ population in the spleen at D9 p.i. of sgCtrl group and sgPdcd1 group.

E. Histogram of PD-1 expression and statistical analysis of PD-1^+^ population from the Cas9^+^sgRNA^+^ P14 cells in the PBMC (D7 p.i.), spleen (D9 p.i.) or liver (D9 p.i.).

F. Sanger sequencing results for the *Pdcd1* locus from FACS sorted Cas9^+^sgRNA^+^ P14 cells from the sgCtrl group (pooled 5 mice) or the sgPdcd1 group (pooled 2 mice).

G. Histogram of KLRG1 or CXCR3 expression and statistical analysis of KLRG1^+^ or CXCR3^+^ populations from Cas9^+^sgRNA^+^ P14 cells from the spleen (D8 p.i. of LCMV-Arm) between targeted-sgRNA and sgCtrl group.

\**P*<0.05, \*\**P*<0.01, \*\*\**P*<0.001, \*\*\*\**P*<0.001 versus control (two-tailed Student’s *t*-test). Data are representative of 2 independent experiments (mean±s.e.m.) with at least 3 mice/group.

**Figure S2 Technical optimization of OpTICS system.**

A-B. Log2 fold change (L2FC) of D8 p.i.(T1) to D2 baseline(T0) across different sgRNAs from spleen (A) or liver (B) under 3 conditions during LCMV-Arm infection. X axis represents different sgRNAs, y axis represents L2FC of D8 p.i.(T1) to D2 baseline(T0). Condition 1: No optimized sorting, average input coverage -- 100 cells/sgRNA, Cas9^+/+^ P14 donor. Condition 2: Optimized sorting (in Star Methods), average input coverage -- 400 cells/sgRNA, Cas9^+/+^ P14 donor. Condition 3: Optimized sorting, average input coverage -- 400 cells/sgRNA, Cas9^+/-^ P14 donor. Example target genes are highlighted with the indicated color.

C. Rank correlation of targeted genes between 2 independent screenings of Cas9^+/+^ or Cas9^+/-^ donor P14 groups. The mean values of sgRNA L2FC from each targeted gene were calculated and ranked from the independent screenings. Pearson correlation of the rankings was calculated.

D. Fold change enrichment of sgPdcd1 at D14 p.i. of LCMV-Cl13 in the spleen. Data from the screening performed in Figure 1A-1C.

E. Pearson correlation of different samples from the screening performed in Figure 1A-1C. Sample collection time (days p.i.), LCMV infection and organ of sorted Cas9^+^sgRNA^+^ cells are presented.

**Figure S3 Genetic deletion of Fli1 leads to a greater T cell expansion.**

A. TIDE assay results showing genome disruption efficiency of *Fli1* locus on the Cas9^+^*Fli1*-sgRNA (sgFli1)^+^ cells. Genome disruption was detected by sanger sequencing.

B. Flow cytometry plots (gated on donor P14 cells) of Cas9 (GFP)^+^sgRNA(VEX)^+^ population on D2 after *in vitro* transduction; D8 and D15 p.i. of Arm from splenocytes from sgCtrl or 2 *Fli1*-sgRNA(sgFli1_290 and sgFli1_360) groups. 5 × 10^4^ activated donor cells were adoptively transferred into infected recipient mice on D1 p.i.

C. Normalized Cas9^+^sgRNA^+^ cell numbers from PBMC, liver and lung from sgCtrl and the two sgFli1 groups at D8 and D15 p.i. of Arm. Cell numbers normalized to the sgCtrl group based on D2 *in vitro* transduction efficiency.

D. Flow cytometry plots (gated on donor P14 cells) of Cas9^+^sgRNA^+^ P14 cells on D2 post *in vitro* transduction, D9 and D15 p.i. with Cl13 (speen) for sgCtrl and the two sgFli1 groups. 5 × 10^4^ activated donor P14 cells were adoptively transferred into the infected recipient mice on D1 p.i.

E. Normalized Cas9^+^sgRNA^+^ cell numbers from PBMC, liver and lung from sgCtrl and the two sgFli1 groups at D9 and D15 p.i. of Cl13. Cell number normalized to the sgCtrl group based on D2 *in vitro* transduction efficiency.

\**P*<0.05, \*\**P*<0.01, \*\*\**P*<0.001, \*\*\*\**P*<0.001 versus control (One-Way Anova analysis). Data are representative of 2-4 independent experiments (mean±s.e.m.) with at least 3 mice/group.

**Figure S4 Fli1 inhibits T**_**EFF**_**-like differentiation on transcriptional and epigenetic levels.**

A-B. Statistical analysis of Ly108^-^CD39^+^ or TCF-1^-^Gzmb^+^ T_EFF_-like cells and Ly108^+^CD39^-^ or TCF-1^+^Gzmb^-^ T_EX_ precursor cell numbers from spleen for sgCtrl and the two sgFli1 groups at D8 (A) and D15 (B) p.i. with Cl13. Gated on Cas9(GFP)^+^ sgRNA (VEX)^+^ P14 cells.

C. Flow cytometry plots and statistical analysis of cell number of CD45.2^+^VEX^+^ P14 for Empty-RV and Fli1-OE-RV groups at D8 and D16 p.i. of Cl13. At D0, CD45.2^+^ P14 cells were activated and CD45.1^+^ recipient mice were infected with Cl13. On D1 p.i., activated P14 were transduced with either Empty-RV or Fli1-OE-RV for 6 hours. On D2 p.i., VEX^+^ P14 cells were sorted from each RV transduced group and 1×10^5^ cells were adoptively transferred into the infected recipients.

D-E. Flow cytometry plots and statistical analysis of Ly108^-^CD39^+^ or TCF-1^-^Gzmb^+^ T_EFF_-like cells and Ly108^+^CD39^-^ or TCF-1^+^Gzmb^-^ T_EX_ precursor frequencies for Empty-RV and Fli1-OE-RV groups on D8 and D16 p.i. of Cl13. Gated on VEX^+^ P14 cells.

F. Statistical analysis of CX3CR1^+^ and Tim-3^+^ frequencies for Empty-RV and Fli1-OE-RV groups on D8 and D16 p.i. of Cl13. Gated on VEX^+^ P14 cells.

G. PCA plot for RNA-seq results of sgCtrl, sgFli1_290 and sgFli1_360 groups on D8 p.i. of Cl13.

H. Overlap of all CUT&RUN Fli1 binding peaks with ATAC-seq detected peaks in sgCtrl and sgFli1 groups.

I. Histogram of all CUT&RUN peaks co-localized with the ATAC-seq peaks. Peaks co-localized with the ATAC-seq peaks are red; peaks not co-localized are blue.

J. CUT&RUN IgG or Fli1 binding signals for P14 cells on D9 p.i. and open chromatin region signals detected by ATAC-seq for sgCtrl, sgFli1_290 and sgFli1_360 groups in the *Tcf7* and *Id3* loci.

\**P*<0.05, \*\**P*<0.01, \*\*\**P*<0.001 versus control (One-Way Anova analysis). Data are representative of 3 independent experiments (mean±s.e.m.) with at least 4 mice/group for A-D.

**Figure S5. Fli1 coordinates with Runx1 and antagonizes Runx3 function.**

A. Flow cytometry plots and statistical analysis of CD45.1^+^mCherry^+^ and Ly108^-^CD39^+^/ Ly108^+^CD39^-^ P14 cell numbers for Empty-RV or Runx1-OE-RV on D2 *in vitro* and D7 p.i. At D0, CD45.1^+^ P14 cells were activated and CD45.2^+^ recipient mice were infected with Cl13; At D1 p.i., activated P14 cells were transduced with Empty-RV or Runx1-OE-RV for 6 hours, and 1×10^5^ transduced P14 cells were adoptively transferred into infected recipient mice. Flow cytometry plots gated on the CD45.1^+^ P14 cells.

B. D2 *in vitro* transduction efficiency of sgCtrl-VEX+Empty-mCherry, sgCtrl-VEX+Runx1-mCherry, sgFli1_290-VEX+Empty-mCherry, and sgFli1_290-VEX+Runx1-mCherry C9P14 cells.

C. Statistical analysis of VEX^+^mCherry^+^ C9P14 cells and Ly108^-^CD39^+^/Ly108^+^CD39^-^ cell numbers from spleen for sgCtrl-VEX+Empty-mCherry, sgCtrl-VEX+Runx1-mCherry, sgFli1_290-VEX+Empty-mCherry, and sgFli1_290-VEX+Runx1-mCherry groups on D7 p.i. of Cl13. Gated on Cas9(GFP)^+^CD45.2^+^ P14 cells.

D. D2 *in vitro* transduction efficiency of sgCtrl-mCherry+Empty-VEX, sgCtrl-mCherry+Runx3-VEX, sgFli1_290-mCherry+Empty-VEX, and sgFli1_290-mCherry+Runx3-VEX C9P14 cells.

E. Statistical analysis of VEX^+^mCherry^+^ C9P14 cells and Ly108^-^CD39^+^/Ly108^+^CD39^-^ cell numbers from spleen for sgCtrl-mCherry+Empty-VEX, sgCtrl-mCherry+Runx3-VEX, sgFli1_290-mCherry+Empty-VEX and sgFli1_290-mCherry+Runx3-VEX at D8 p.i. of Cl13. Gated on Cas9^+^CD45.2^+^ P14 cells.

\**P*<0.05, \*\**P*<0.01 versus control (two-tailed Student’s *t*-test). Data are representative of 2 independent experiments (mean±s.e.m.) with at least 5 mice/group.

**Figure S6. *Fli1*-deficiency results in CD8 T cell expansion during influenza virus or *Listeria monocytogenes* infection.**

A. Flow cytometry plots showing sgRNA(VEX)^+^ C9P14 cells in the lungs of influenza virus (PR8-GP33) infected mice, comparing non-transfer(NT), sgCtrl and sgFli1 groups at D8 p.i. “Recovered” was defined by complete weight recovery at D8 p.i.

B. Correlation of sgRNA^+^ C9P14 cell numbers and weight ratio (D8 p.i./D2 p.i.) during the PR8-GP33 infection. In the “weight not recovered” group (weight ratio between 0.7 and 1.0), sgRNA^+^ C9P14 cell numbers were further compared.

C. Flow cytometry plots and statistical analysis of sgRNA^+^ C9P14 cells in the spleens of PR8-GP33-infected recipient mice for non-transfer (NT), sgCtrl and sgFli1 groups at D8 p.i.

D. Flow cytometry plots and statistical analysis of sgRNA^+^ C9P14 cells in the spleens of *Listeria monocytogenes*-infected recipients for non-transfer (NT), sgCtrl and sgFli1 groups at D7 p.i.

\**P*<0.05, \*\**P*<0.01 versus control (two-tailed Student’s *t*-test and One-Way Anova analysis). Data are representative of 2 independent experiments (mean±s.e.m.) with at least 6 mice/group.

**Figure S7. Deleting Fli1 in CD8 T cells leads to better tumor protection in an immune competent setting.**

A. Experimental design. On D0, CD45.2^+^Cas9^+^P14^-^ mice were inoculated with 2 x 10^5^ B16-D^b^gp33 cells. On D3 post tumor inoculation (p.t.), CD8 T cells were isolated from CD45.2^+^ C9P14 donor mice and activated. The next day, activated C9P14 cells were transduced with sgCtrl or sgFli1 RV for 6 hours. On D5 p.t., sgRNA (VEX)^+^ P14 cells were sorted from sgCtrl or sgFli1 groups, and 3 x 10^6^ purified VEX^+^ C9P14 cells were adoptively transferred into tumor-bearing mice.

B. Tumor volume curve of tumor-bearing mice from NT, sgCtrl^+^ and sgFli1^+^ C9P14 cell transferred groups.

C. Tumor weight from NT, sgCtrl^+^ and sgFli1^+^ C9P14 transferred mice on D24 p.t.

D-E. Flow cytometry plots (D) and statistical analysis (E) of sgRNA (VEX)^+^ C9P14 cells and Ly108^-^ CD39^+^/Ly108^+^CD39^-^ populations from tumor for sgCtrl and sgFli1 groups on D24 p.t.

F-G. Statistical analysis of sgRNA^+^ C9P14 cells and Ly108^-^CD39^+^/Ly108^+^CD39^-^ populations from draining lymph node (dLN, F) and spleen (G) for sgCtrl and sgFli1 groups on D24 p.t.

\**P*<0.05, \*\**P*<0.01, \*\*\**P*<0.001, \*\*\*\**P*<0.001 versus control (two-tailed Student’s *t*-test and One-Way Anova analysis). Data are representative of 3 independent experiments (mean±s.e.m.) with at least 6 mice/group.

