## Supplementary figures for "*In vivo* CRISPR screening identifies Fli1 as a transcriptional safeguard that restrains effector CD8 T cell differentiation during infection and cancer"

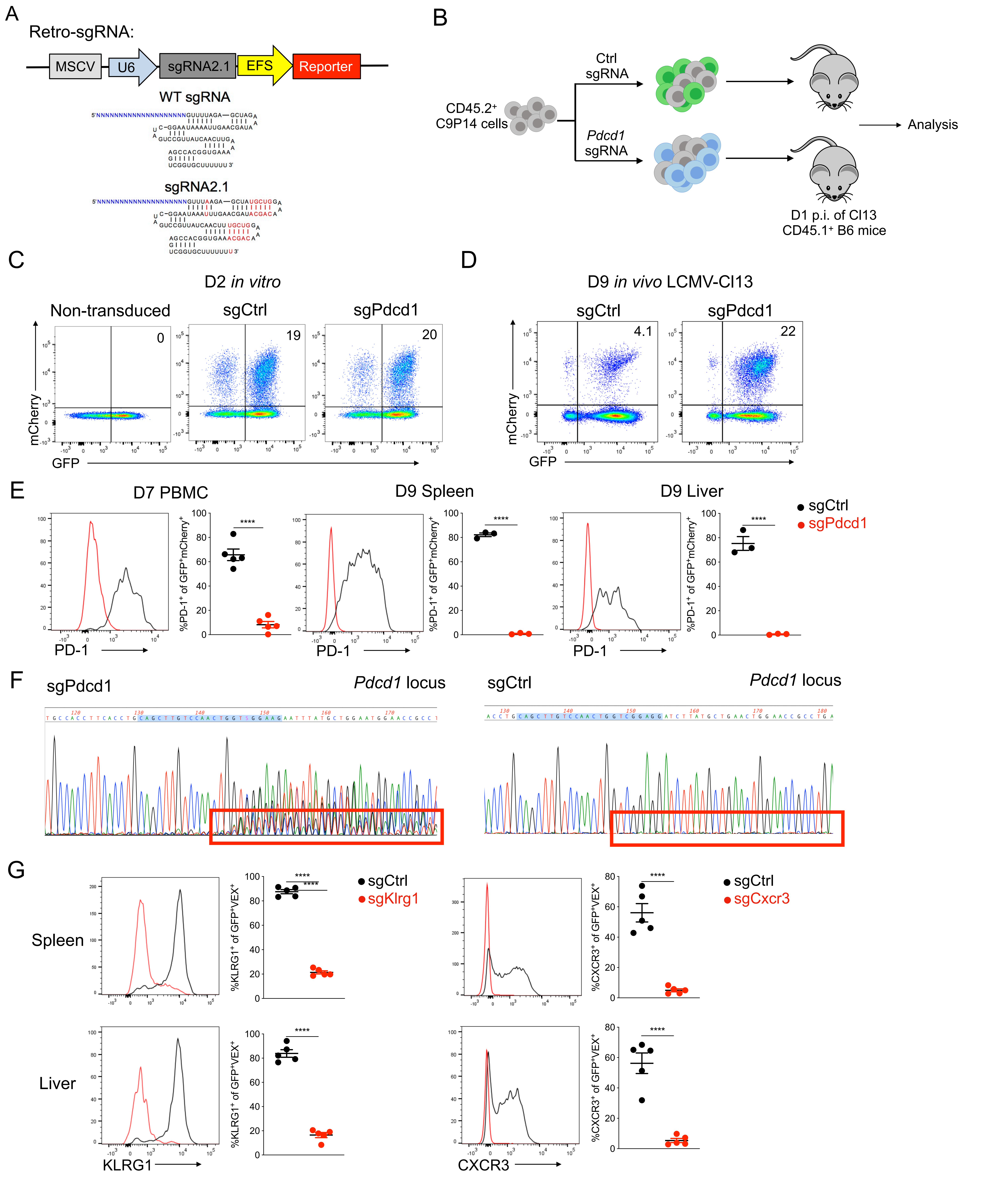
A
Retro-sgRNA:
MSCV
U6
sgRNA2.1
EFS
Reporter
WT sgRNA
sgRNA2.1
B
CD45.2⁺ C9P14 cells
Ctrl sgRNA
Pdcd1 sgRNA
Analysis
D1 p.i. of Cl13 CD45.1⁺ B6 mice
C
D2 in vitro
Non-transduced
sgCtrl
sgPdcd1
0
19
20
mCherry
GFP
D
D9 in vivo LCMV-Cl13
sgCtrl
sgPdcd1
4.1
22
mCherry
GFP
E
D7 PBMC
D9 Spleen
D9 Liver
sgCtrl
sgPdcd1
%PD-1⁺ of GFP⁺mCherry⁺
PD-1
****
F
sgPdcd1
Pdcd1 locus
sgCtrl
Pdcd1 locus
G
Spleen
Liver
sgCtrl
sgKlrg1
sgCxcr3
%KLRG1⁺ of GFP⁺VEX⁺
%CXCR3⁺ of GFP⁺VEX⁺
KLRG1
CXCR3
****

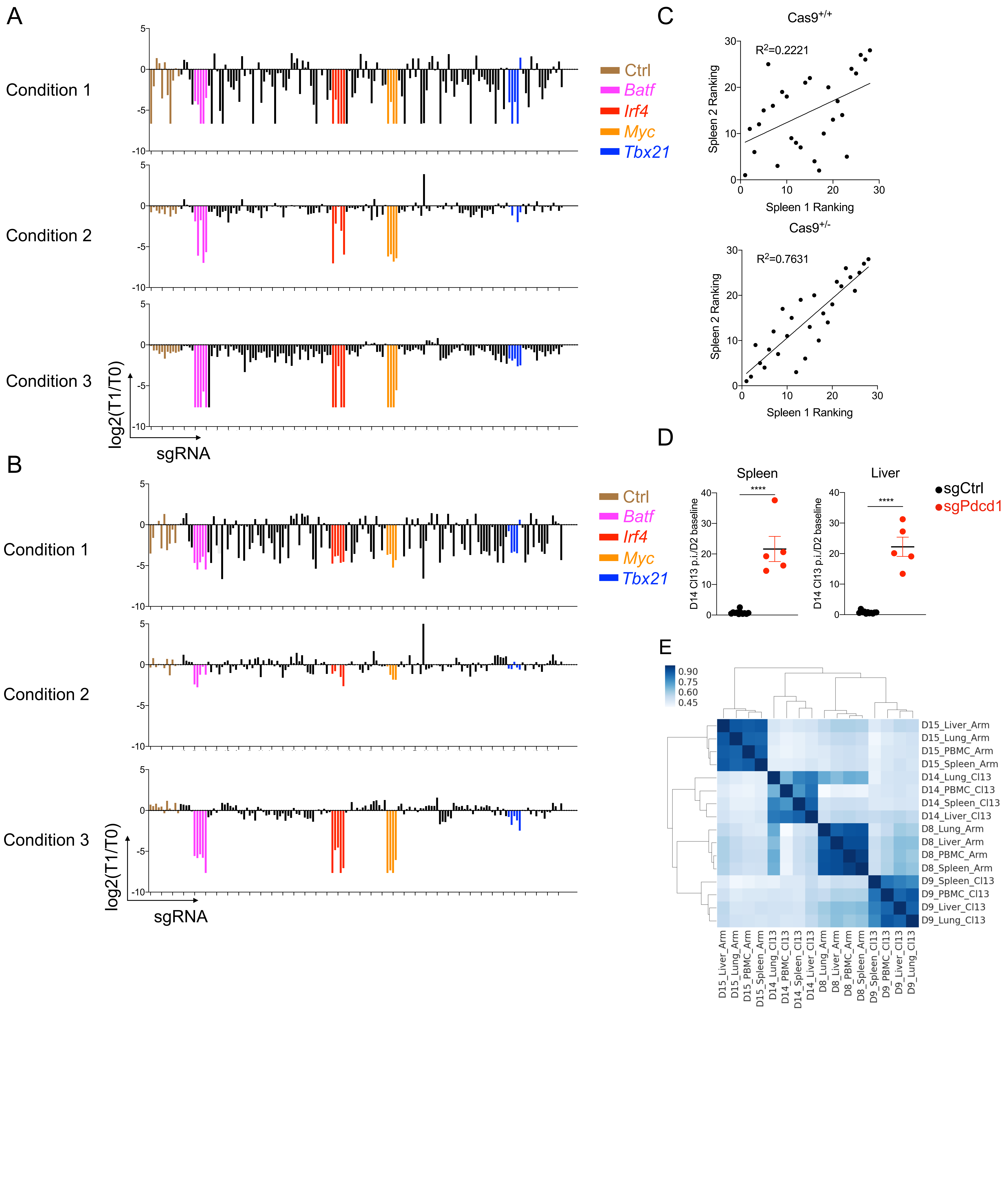


Figure S2, related to Figure 1

A

| Sample Name | TIDE Efficiency(%) |
| --- | --- |
| D8 Arm Fli1_290_1 | 76.4 |
| D8 Arm Fli1_290_2 | 72.6 |
| D8 Arm Fli1_360_1 | 69.2 |
| D8 Arm Fli1_360_2 | 71.7 |
| D9 Cl13 Fli1_290 | 80.8 |
| D9 Cl13 Fli1_360 | 79.8 |

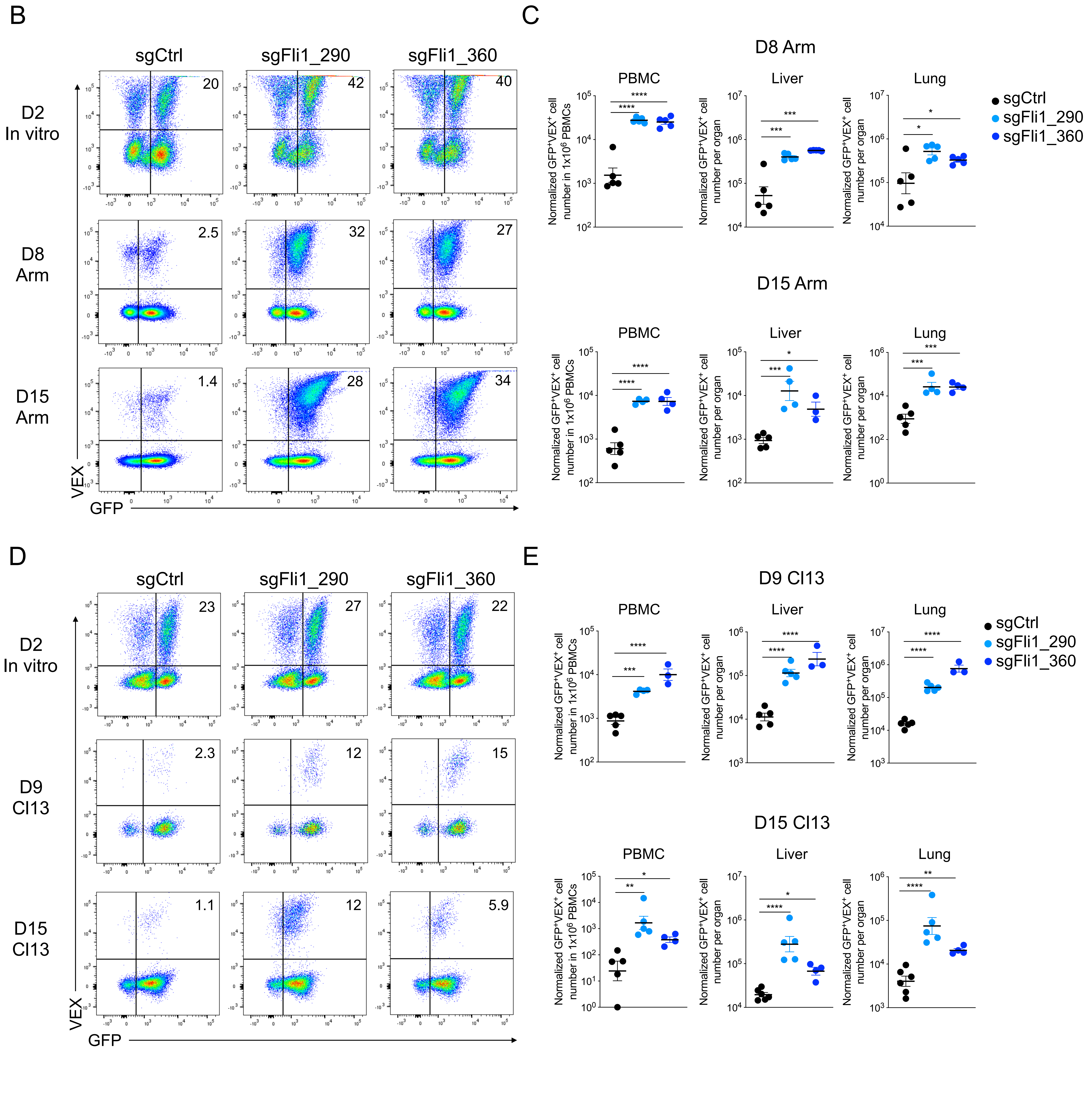


Figure S3, related to Figure 1

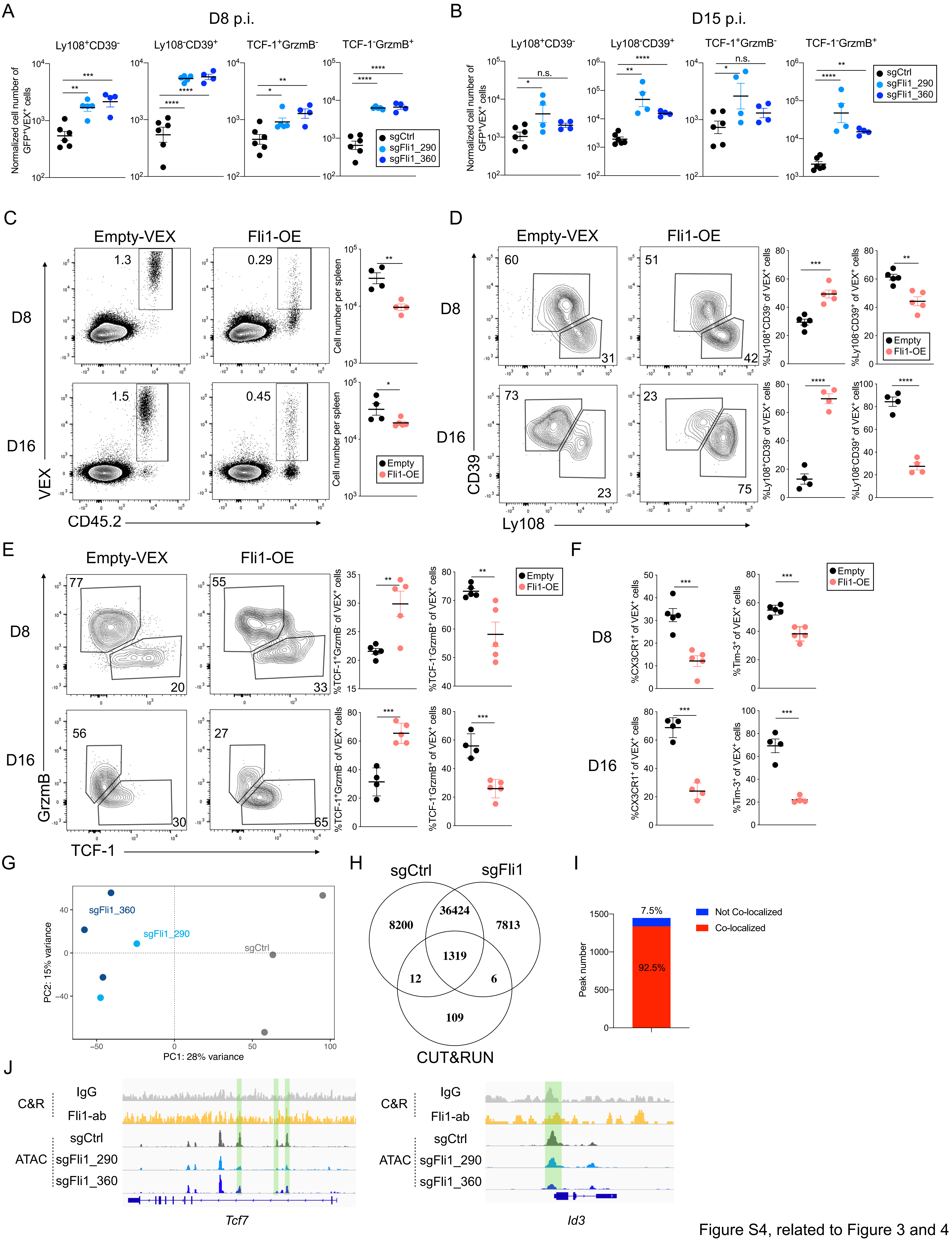


Figure S4, related to Figure 3 and 4

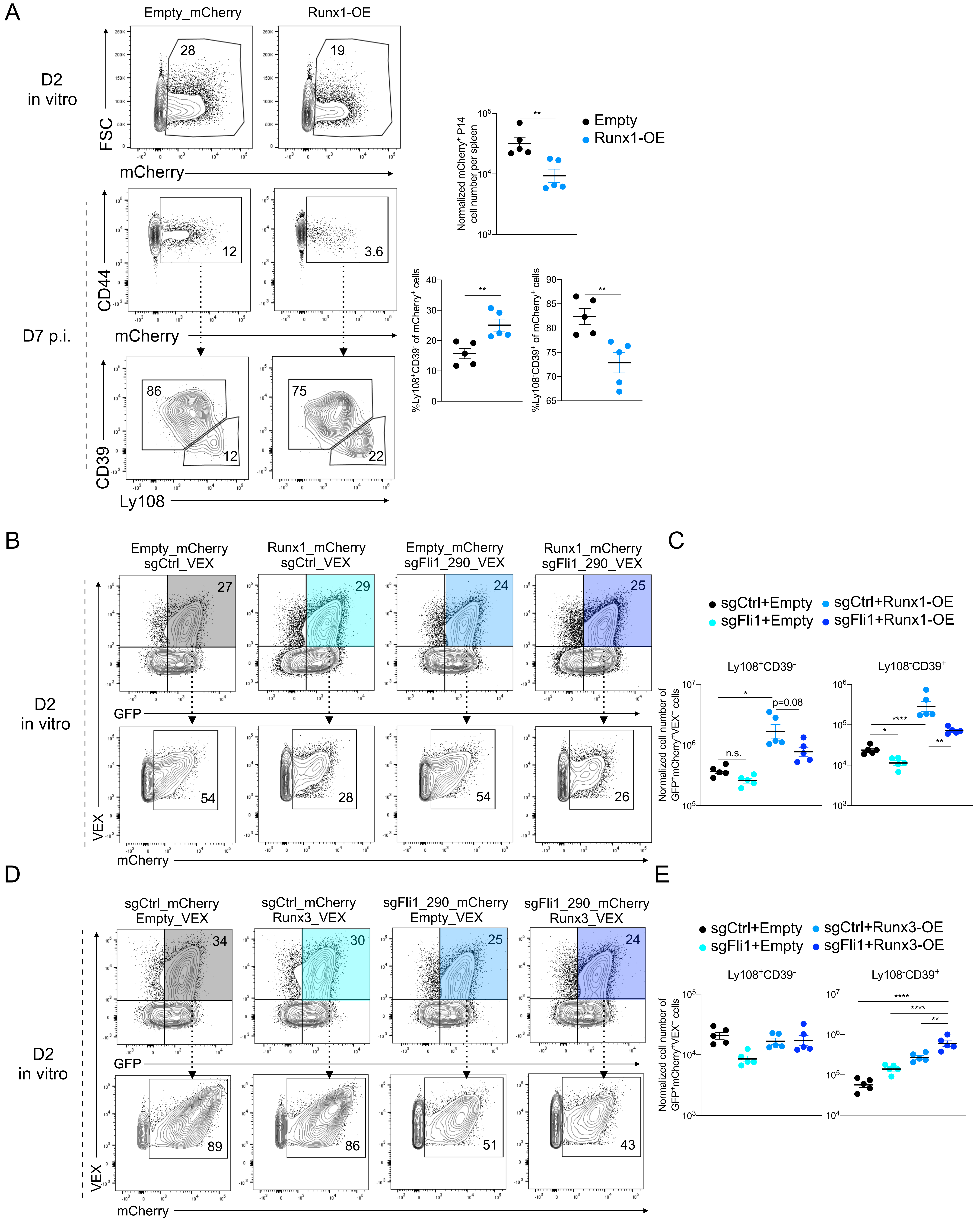


Figure S5, related to Figure 5

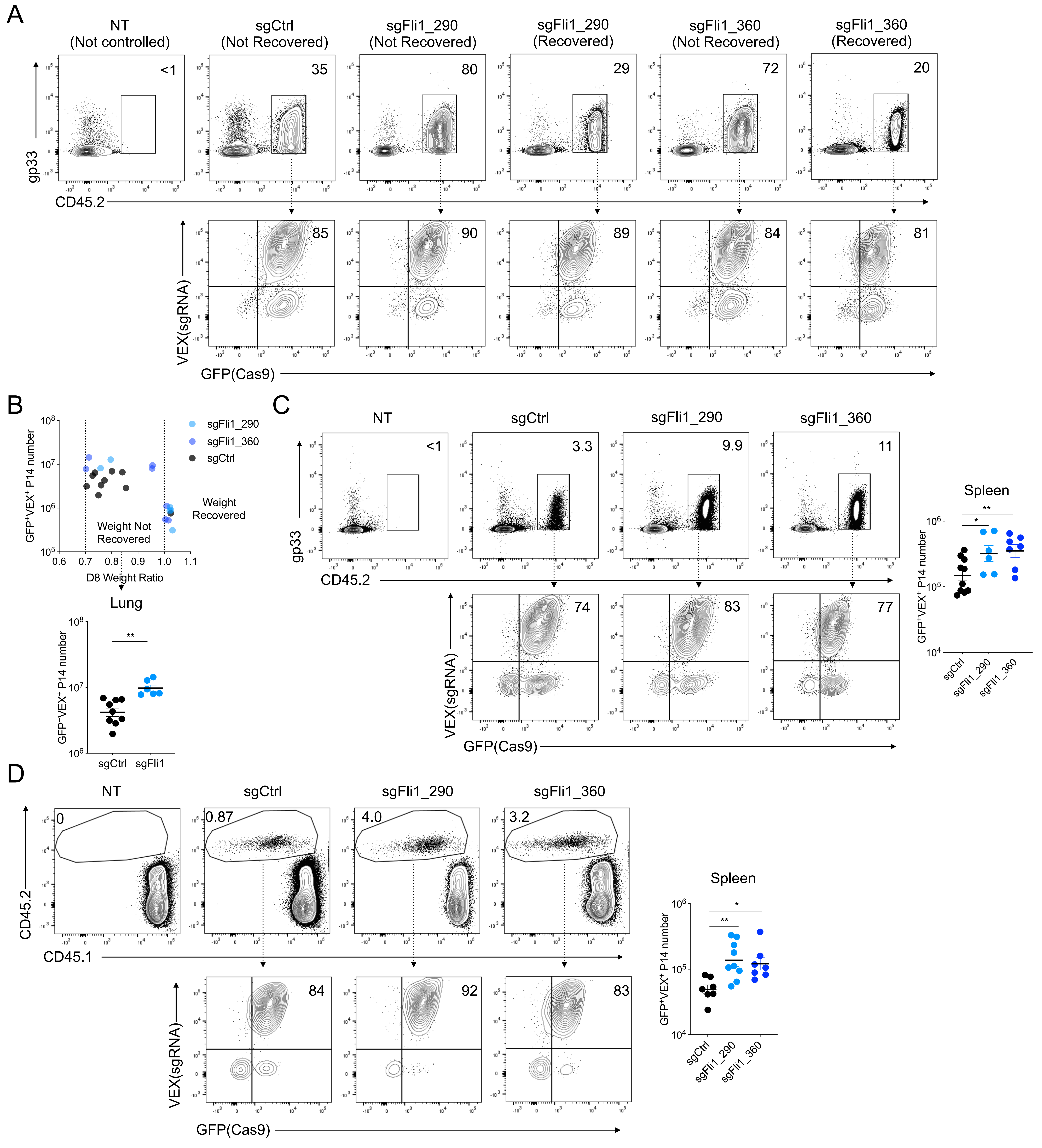


Figure S6 related to Figure 6

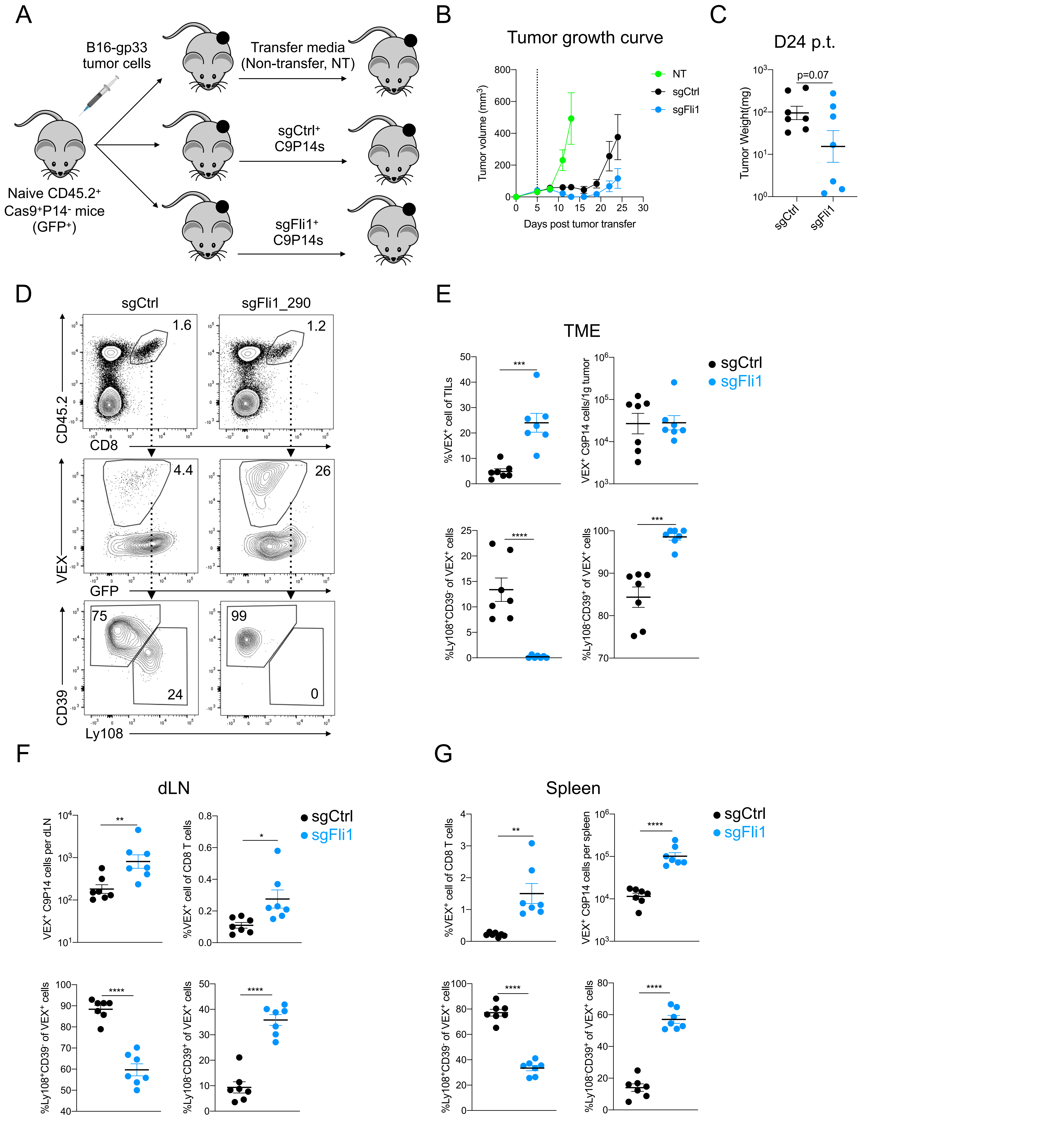


Figure S7 related to Figure 7
